# Environmental and genetic drivers of population differences in SARS-CoV-2 immune responses

**DOI:** 10.1101/2022.11.22.517073

**Authors:** Yann Aquino, Aurélie Bisiaux, Zhi Li, Mary O’Neill, Javier Mendoza-Revilla, Sarah Hélène Merkling, Gaspard Kerner, Milena Hasan, Valentina Libri, Vincent Bondet, Nikaïa Smith, Camille de Cevins, Mickaël Ménager, Francesca Luca, Roger Pique-Regi, Giovanna Barba-Spaeth, Stefano Pietropaoli, Olivier Schwartz, Geert Leroux-Roels, Cheuk-Kwong Lee, Kathy Leung, Joseph T.K. Wu, Malik Peiris, Roberto Bruzzone, Laurent Abel, Jean-Laurent Casanova, Sophie A. Valkenburg, Darragh Duffy, Etienne Patin, Maxime Rotival, Lluis Quintana-Murci

## Abstract

Humans display vast clinical variability upon SARS-CoV-2 infection^1–3^, partly due to genetic and immunological factors^4^. However, the magnitude of population differences in immune responses to SARS-CoV-2 and the mechanisms underlying such variation remain unknown. Here we report single-cell RNA-sequencing data for peripheral blood mononuclear cells from 222 healthy donors of various ancestries stimulated with SARS-CoV-2 or influenza A virus. We show that SARS-CoV-2 induces a weaker, but more heterogeneous interferon-stimulated gene activity than influenza A virus, and a unique pro-inflammatory signature in myeloid cells. We observe marked population differences in transcriptional responses to viral exposure that reflect environmentally induced cellular heterogeneity, as illustrated by higher rates of cytomegalovirus infection, affecting lymphoid cells, in African-descent individuals. Expression quantitative trait loci and mediation analyses reveal a broad effect of cell proportions on population differences in immune responses, with genetic variants having a narrower but stronger effect on specific loci. Additionally, natural selection has increased immune response differentiation across populations, particularly for variants associated with SARS-CoV-2 responses in East Asians. We document the cellular and molecular mechanisms through which Neanderthal introgression has altered immune functions, such as its impact on the myeloid response in Europeans. Finally, colocalization analyses reveal an overlap between the genetic architecture of immune responses to SARS-CoV-2 and COVID-19 severity. Collectively, these findings suggest that adaptive evolution targeting immunity has also contributed to current disparities in COVID-19 risk.

## Introduction

One of the most striking features of the COVID-19 pandemic is the remarkable extent of clinical variation among SARS-CoV-2 infected individuals, ranging from asymptomatic infection to lethal disease^1–3^. Risk factors include primarily advanced age^1^ but also male sex^5^, comorbidities^6^, and human genetic factors (i.e., rare and common variants)^4,7^. Furthermore, variation in innate immunity^8–10^ – including inborn errors or neutralizing auto-antibodies against type I interferons^11–13^ – contribute to the various SARS-CoV-2-related clinical manifestations, and epidemiological and genetic data suggest differences in the outcome of SARS-CoV-2 infection between populations^6,7,14,15^. These observations, together with previous reports on the importance of ancestry-related differences in transcriptional responses to immune challenges^16–19^, highlight the need for in-depth investigations of the magnitude of variation in immune responses to SARS-CoV-2 and its drivers across populations worldwide.

There is strong evidence to suggest that viruses and other infectious agents have had an overwhelming impact on human evolution, exerting selection pressures that have shaped present-day population genetic variation^20^. In particular, human adaptation to RNA viruses, through selective sweeps or admixture with archaic hominins, has been identified as a source of population genetic differentiation^17,21,22^. For example, strong genetic adaptation, starting ~25,000 years ago, has targeted multiple human coronavirus-interacting proteins in East Asian populations^23,24^. Furthermore, there is growing evidence for links between Neanderthal introgression and immunity^25^, with reports of COVID-19 severity being modulated by Neanderthal haplotypes in modern Eurasians^26,27^. However, the ways in which past natural selection events and archaic admixture have affected the immune response to SARS-CoV-2 in contemporary humans remains to be investigated.

We addressed these questions by exposing peripheral blood mononuclear cells (PBMCs) from individuals of Central African, West European, and East Asian descent to SARS-CoV-2 and, for the purpose of comparison, to another respiratory RNA virus, influenza A virus (IAV). By combining single-cell RNA sequencing (scRNA-seq) data with quantitative and population genetics approaches, we delineate the respective contributions of cellular, genetic, and evolutionary factors to population variation in immune responses to SARS-CoV-2.

### Defining single-cell responses to RNA viruses

We characterized cell type-specific transcriptional responses to SARS-CoV-2 and IAV, by performing scRNA-seq on PBMCs from 222 healthy donors who originate from three different geographic locations (Central Africa, *n* = 80; West Europe, *n* = 80; East Asia, *n* = 62; Methods), thus exposed to probably different environmental conditions, and carry different genetic ancestries (Supplementary Fig. 1). PBMCs were treated for six hours (i.e., a time point at which immune responses were strong and cell viability was high; Supplementary Fig. 2, Supplementary Note 1, Supplementary Table 1) with a mock-control (non-stimulated), SARS-CoV-2 (ancestral strain, BetaCoV/France/GE1973/2020) or IAV (H1N1/PR/8/1934) (*n=222* samples for each set of experimental conditions). We captured over one million high-quality single-cell transcriptomes (Fig. 1a, Supplementary Fig. 3, Supplementary Table 2a). By combining transcriptome-based clusters with cellular indexing of transcriptomes and epitopes by sequencing on a subset of samples (CITE-seq; Methods), we defined 22 different cell types across five major immune lineages, including myeloid cells, B cells, CD4^+^ T cells, CD8^+^ T cells and natural killer (NK) cells (Fig. 1b, Supplementary Fig. 4, Supplementary Table 2b-d).

**Figure 1.**
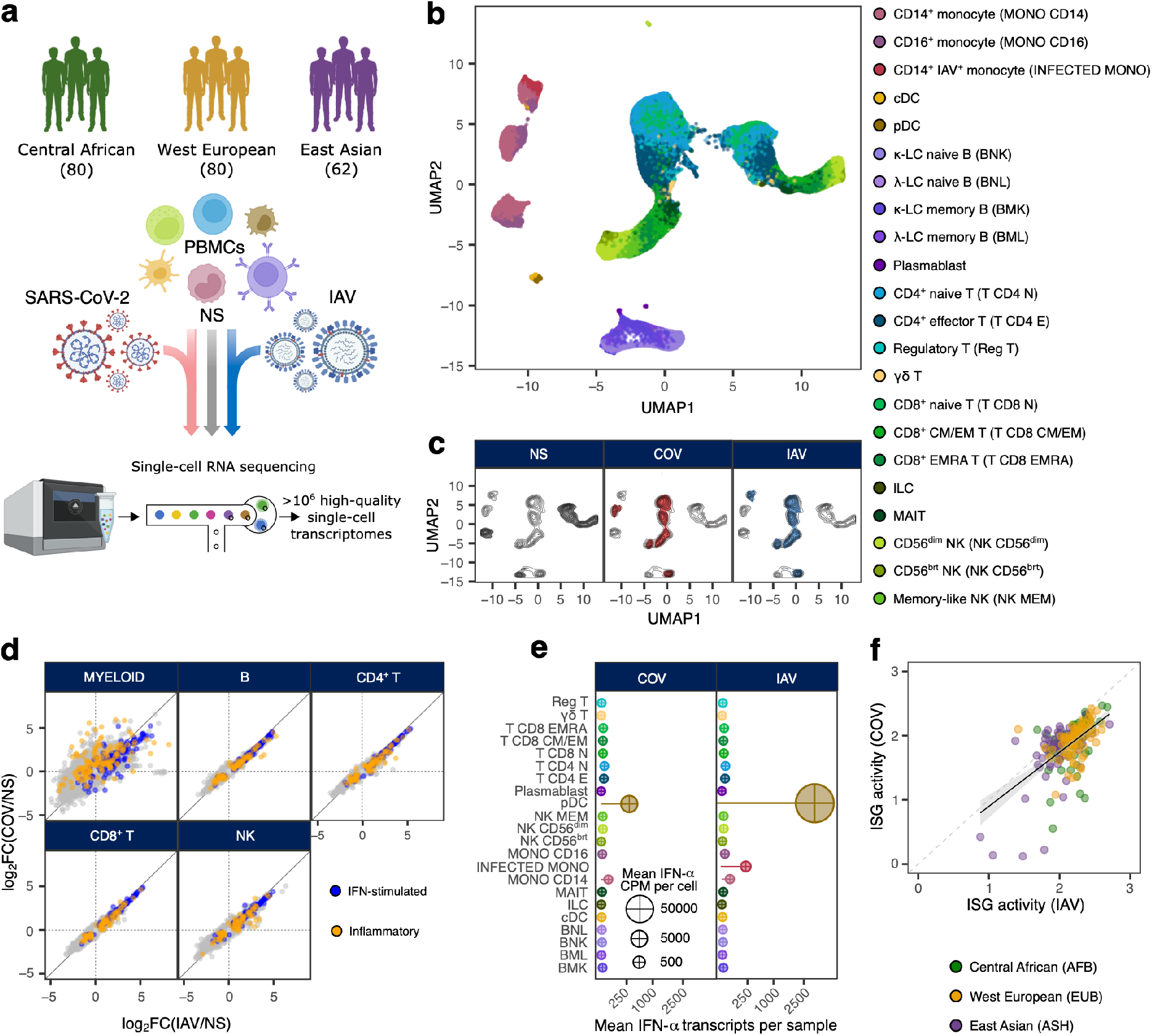
Population-scale single-cell responses to SARS-CoV-2 and IAV. **a,** Study design. **b** and **c,** Uniform manifold approximation and projection (UMAP) of 1,047,824 peripheral blood mononuclear cells: resting (non-stimulated; NS), stimulated with SARS-CoV-2 (COV), or influenza A virus (IAV) for six hours. **b**, The colors indicate the 22 different cell types inferred. **c**, Distribution of cells in NS, COV and IAV conditions on UMAP coordinates. Contour plot indicates the overall density of cells, and colored areas delineate regions of high cell density in each condition (gray: NS, red: COV, blue: IAV). **d**, Comparison of transcriptional responses to SARS-CoV-2 and IAV across major immune lineages. Hallmark inflammatory and interferon-stimulated genes are highlighted in orange and blue, respectively. **e**, Relative expression of IFN-α-encoding transcripts by each immune cell type in response to SARS-CoV-2 and IAV. Bar lengths indicate the mean number of IFN-α transcripts contributed by each cell type to the overall pool (cell type frequency × mean number of IFN-α transcripts per cell). Dot area is proportional to the mean level of IFN-α transcripts in each cell type (counts per million). **f**, Correlation of ISG activity scores between individuals, following exposure to SARS-CoV-2 and IAV. Each dot corresponds to a single individual (*n* = 222) and its color indicates the self-reported ancestry of the individual concerned (AFB: Central African; EUB: West European; ASH: East Asian).

After adjusting for technical factors (Methods), we found that cell-type identity was the main driver of gene expression variation (~32%), followed by virus exposure (~27%) (Fig. 1b, c, Supplementary Fig. 5). Both SARS-CoV-2 and IAV induced a strong immune response, with 2,914 genes upregulated (FDR < 0.01, log_2_FC > 0.5; out of 12,655 with detectable expression) in response to virus stimulation across cell lineages (Supplementary Table 2e). Transcriptional responses to these viruses were highly correlated across cell types and were characterized by a strong induction of interferon-stimulated genes (ISG) (Fig. 1d). However, we observed marked heterogeneity in the myeloid response, with SARS-CoV-2 inducing a specific transcriptional network enriched in inflammatory-response genes (GO:0006954; fold-enrichment (FE) = 3.4, FDR < 4.9 × 10^-8^) (Supplementary Table 2f). For example, *IL1A, IL1B* and *CXCL8*, encoding pro-inflammatory cytokines, were highly and specifically upregulated in response to SARS-CoV-2 (log_2_FC > 2.8, FDR < 2.3 × 10^-36^), highlighting the greater inflammatory potential of this virus.

We assessed interindividual variability in the response to viral stimuli, by summarizing the response of each individual as a function of their mean level of ISG expression (i.e., ISG activity; Methods, Supplementary Table 2g). We found that SARS-CoV-2 induced more variable ISG activity than IAV across lineages^28^, with myeloid cells displaying the strongest differences (Supplementary Fig. 6a). We determined the relative contributions of the various interferons (IFNs) to the variation of ISG activity, by using single-molecule arrays (SIMOA) to quantify the levels of secreted IFN-α, β and γ proteins. In the SARS-CoV-2 condition, IFN-α alone accounted for up to 57% of ISG variability, highlighting its determinant role in the response to SARS-CoV-2 (Supplementary Fig. 6b, c). IFN-α transcripts were produced by both infected CD14^+^ monocytes and plasmacytoid dendritic cells (pDCs) after stimulation with IAV, but pDCs were the only source of IFN-α after stimulation with SARS-CoV-2 (Fig. 1e) and these cells presented lower levels of *IFNA1-21* expression (log_2_FC = 6.4 for SARS-CoV-2 *vs*. 12.5 for IAV, Wilcoxon rank-sum *p*-value = 1.2 × 10^-16^). Nevertheless, patterns of interindividual variability for ISG activity were remarkably similar after treatment with SARS-CoV-2 and IAV (*r* = 0.60, Pearson’s *p*-value < 1.2 × 10^-22^, Fig. 1f), indicating that, at the population level, the IFN-driven response is largely shared between these two viruses.

### Marked cellular heterogeneity across populations

We investigated the contribution of differences in cellular proportions to the observed interindividual variability of SARS-CoV-2 responses, by focusing on individuals of Central African and West European ancestries — all recruited during the same sampling campaign, thereby mitigating any potential batch effects related to sample processing^17^. We detected marked differences in lineage composition across populations, particularly for NK cells (Fig. 2a, Supplementary Table 3a). Notably, an NK subset, identified as memory-like NK cells^29^, constituted 55.2% of the NK compartment of African-ancestry individuals, but only 12.2% of European-ancestry individuals (Wilcoxon rank-sum *p*-value < 6.4 × 10^-20^; Supplementary Fig. 7a, b). Individuals of African ancestry also presented higher proportions of CD16^+^ monocytes^19^ and memory lymphocyte subsets, such as memory B cells, effector CD4^+^ T cells and effector memory CD8^+^ T cells re-expressing CD45RA (CD8^+^ EMRA T cells) (Wilcoxon rank-sum *p*-value < 4.1 × 10^-5^).

**Figure 2.**
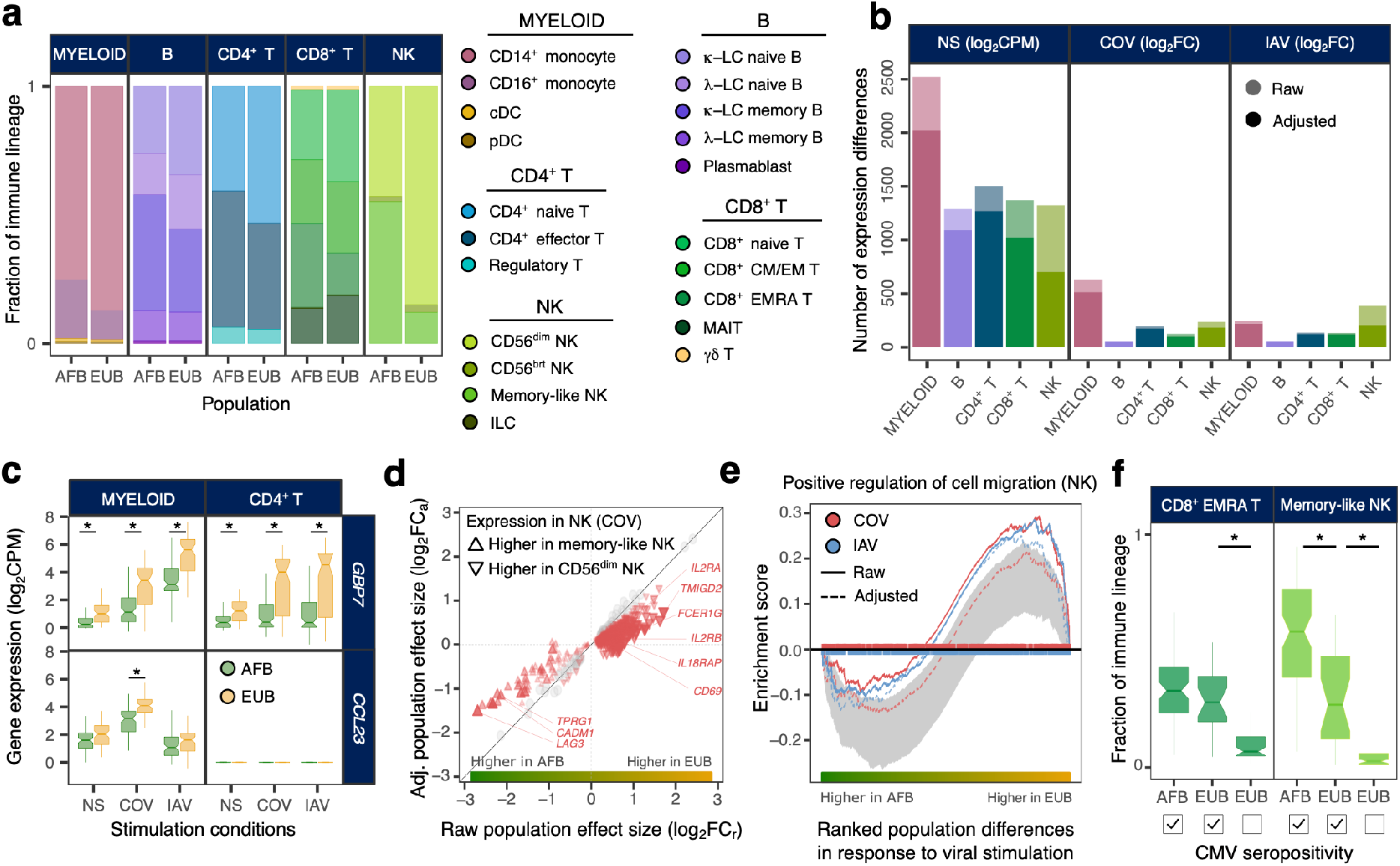
Effects of the variation of cellular composition on transcriptional responses to viral stimuli. **a,** Cell-type proportions within each major immune lineage in individuals of Central African (AFB) and West European (EUB) ancestries. **b,** Number of genes differentially expressed between AFB and EUB donors, in the basal state (NS) or in response to SARS-CoV-2 (COV) or influenza A virus (IAV), in each major immune lineage. Numbers are provided before and after adjustment for cellular composition. **c**, Examples of genes displaying population differential responses (popDRGs), either shared between cell types and viruses (*GBP7*) or specific to SARS-CoV-2-stimulated myeloid cells (*CCL23*). *Benjamini-Hochberg adjusted *p*-value < 0.001. **d,** Effect of adjusting for cellular composition on genes differentially expressed between populations. Adjustment reduces the number of genes with different expression levels between memory-like NK cells and their non-memory counterpart (i.e., CD56^dim^ NK cells) following exposure to SARS-CoV-2 (red triangles). **e,** Effect of adjusting for cellular composition on population differences in the response to viral stimulation for genes involved in the ‘positive regulation of cell migration’ (GO:0030335) in the NK lineage. **f,** Distribution of CD8^+^ EMRA T and memory-like NK cell frequencies in AFB and EUB donors according to cytomegalovirus serostatus (CMV^+/-^) *Wilcoxon Rank-Sum *p*-value < 0.001. **d** and **f**, middle line: median; notches: 95% confidence intervals (CI) of median, box limits: upper and lower quartiles; whiskers: 1.5× interquartile range.

We then searched for genes displaying differences in expression between populations. We found 3,389 genes, across lineages, differentially expressed between populations (popDEGs; FDR < 0.01, |log_2_FC| > 0.2) in the basal state, and 898 and 652 genes displaying differential responses between populations (popDRGs; FDR < 0.01, |log_2_FC| > 0.2) after stimulation with SARS-CoV-2 and IAV, respectively (Fig. 2b, Supplementary Table 3b, c). The popDRGs included genes encoding key immunity regulators, such as the IFN-responsive GBP7 and the macrophage inflammatory protein CCL23 (MIP-3), both of which were more strongly upregulated in Europeans (Fig. 2c). The *GBP7* response was common to both viruses and all lineages (log_2_FC > 0.88, Student’s *t*-test adj. *p*-value < 1.4 × 10^-3^), but that of *CCL23* was specific to SARS-CoV-2-stimulated myeloid cells (log_2_FC = 0.72, Student’s adj. *p*-value = 5.3 × 10^-4^). We estimated that population differences in cellular composition accounted for 15-47% of popDEGs and for 7-46% of popDRGs, with the strongest impact on NK cells (Fig. 2b, d, Supplementary Fig. 7c). The variation of cellular composition mediated pathway-level differences in response to viral stimulation between populations (Supplementary Table 3d). For example, in virus-stimulated NK cells, genes involved in the promotion of cell migration, such as *CSF1* or *CXCL10*, were more strongly induced in donors of European ancestry (normalized enrichment score > 1.5, Gene Set Enrichment Analysis adj. *p*-value < 0.009). However, the loss of this signal after adjustment for cellular composition (Fig. 2e) indicates that fine-scale cellular heterogeneity drives population differences in immune responses to SARS-CoV-2.

### Repercussions of latent cytomegalovirus infection

Latent cytomegalovirus (CMV) infection has been reported to vary across populations worldwide^30^ and to alter cellular proportions^31–33^. We therefore determined the CMV serostatus of the samples. All but one of the individuals of Central African ancestry were CMV^+^ (99%), versus only 31% of donors of West European ancestry, and CMV seropositivity was strongly correlated with the proportions of memory-like NK and CD8^+^ EMRA T cells (Fig. 2f, Supplementary Fig. 7d). Using mediation analysis, we estimated that CMV serostatus accounts for up to 73% of population differences in the proportion of these cell types (Supplementary Table 3e). These differences had a profound impact on the transcriptional response to SARS-CoV-2 (Supplementary Note 2, Supplementary Table 3f), probably contributing to the reported associations between CMV serostatus and COVID-19 severity^34,35^. However, other than its effects on cellular composition, we found that CMV infection had a limited direct effect on the response to SARS-CoV-2, with only one gene presenting significant differences in expression in response to this virus at a FDR of 1% (*ERICH3* in CD8^+^ T cells, log_2_FC = 1.7, FDR = 0.007; Supplementary Table 3g). These results reveal how environmental exposures that differ between populations, such as CMV infection, can lead to changes in the composition of the lymphoid fraction that, in turn, explain the observed population differences in the response to SARS-CoV-2.

### Genetic architecture of the leukocyte response

We assessed the effects of human genetic variants on transcriptional variation, by mapping expression quantitative trait loci (eQTLs), focusing on *cis*-regulatory variants. At a FDR of 1%, we identified between 1,866 and 4,323 independent eQTLs per major cell lineage, affecting a total of 5,198 genes (Fig. 3a, Supplementary Table 4a). Increasing the resolution to 22 cell types led to the identification of an additional 3,603 eQTLs (Fig. 3b, Supplementary Fig. 8a, Supplementary Table 4b), highlighting the value of scRNA-seq for identifying context-dependent eQTLs. We found that 79% of eQTLs were replicated (*p*-value < 0.01) in at least three cell types, but only 22% were common to all lineages. In total, 812 eQTLs were cell type-specific (Methods), ~45% of which were detected in myeloid cells (Fig. 3b), including a pDC-specific eQTL (rs114273142) affecting the host gene encoding miRNA-155 — a miRNA that ultimately promotes sensitivity to type I IFNs^36^ (Supplementary Fig. 8b). More broadly, we found that eQTL effect sizes were highly correlated across ontogenetically related cell types (mean correlation within and between lineages of *r* = 0.60 and 0.47, respectively, Wilcoxon rank-sum *p*-value = 6.2 × 10^-6^; Fig. 3c).

**Figure 3.**
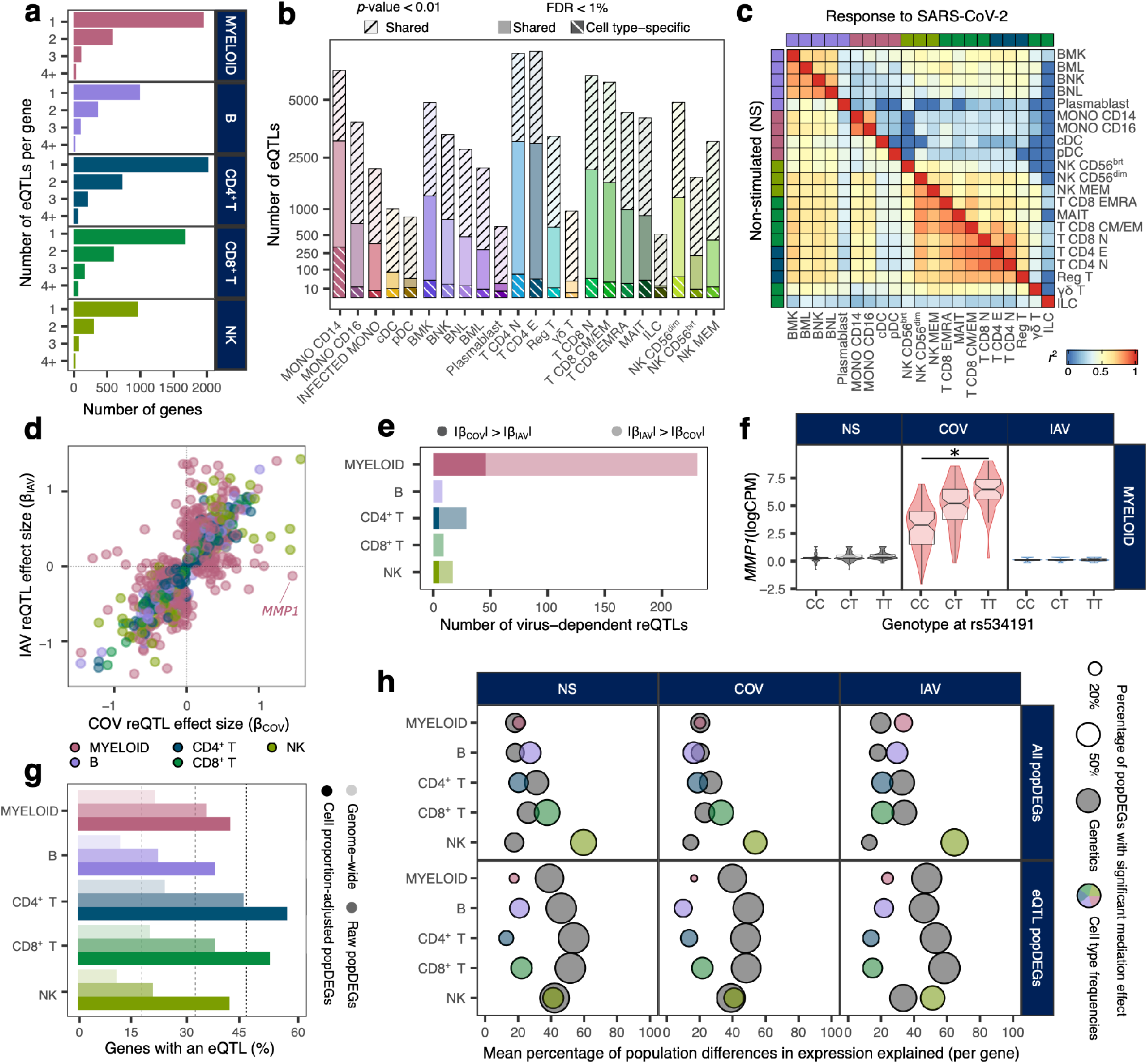
Genetic architecture of the variation of immune response to RNA viruses. **a**, Number of eQTLs detected per gene within each major immune lineage. **b**, Total number of eQTLs detected in each of the 22 different cell types. Colored bars indicate the number of genome-wide significant eQTLs in each cell type, white stripes (bottom) indicate cell type-specific eQTLs (*p*-value > 0.01 in all other cell types), and black stripes (top) indicate the total number of eQTLs detected in each cell type including eQTLs from other cell types replicated at a *p*-value < 0.01. **c**, Correlation of eQTL (NS; lower triangle) and reQTL (response to SARS-CoV-2; upper triangle) effect sizes across cell types. For each pair of cell types, Spearman’s correlation coefficient was calculated for the effect sizes (β) of eQTLs that are significant at a nominal *p*-value < 0.01 in each cell type. **d**, Comparison of reQTL effect sizes (β) between SARS-CoV-2- and IAV-stimulated cells. Each dot represents a specific reQTL (i.e., SNP, gene, and lineage) and its color indicates the immune lineage in which it was detected. **e**. Number of virus-dependent reQTLs (interaction *p*-value < 0.01) in each immune lineage, split according to the stimulus where the reQTL has the largest effect size. **f**, Example of a SARS-CoV-2-specific reQTL at the *MMP1* locus. *Student’s *t*-test *p*-value < 10^-16^; middle line: median; notches: 95% CI of median, box limits: upper and lower quartiles; whiskers: 1.5× interquartile range; points: outliers. **g,** Enrichment in eQTLs among genes differentially expressed between populations (popDEGs). For each immune lineage, bars indicate the percentage of genes with a significant eQTL, at the genome-wide scale and among popDEGs, before or after adjustment for cellular composition. **h**, For each immune lineage and stimulation condition, the *x*-axis indicates the mean percentage difference in expression between populations mediated by genetics (i.e., the most significant eQTL per gene in each immune lineage and condition) or cellular composition, across all popDEGs (top) or the set of popDEGs associated with a significant eQTL (bottom). The size of the dots reflects the percentage of genes with a significant mediated effect at a FDR of 1%.

We then focused on genetic variants that altered the response to viral stimuli (i.e., response eQTLs, reQTLs). We identified 1,505 reQTLs affecting 1,213 genes (Supplementary Table 4c, d). The correlation of effect sizes across ontogenetically related cell types was weaker for reQTLs than for eQTLs (0.36 *vs*. 0.50, respectively, Wilcoxon rank-sum *p*-value < 5.6 × 10^-13^, Fig. 3c). Furthermore, the proportion of shared reQTLs between the two viruses differed between cell types. In lymphoid cells, 93% of the reQTLs detected after stimulation with SARS-CoV-2 were also detected after stimulation with IAV (*p*-value < 0.01), with only 7.7% differing in effect size between viruses (interaction *p*-value < 0.01; Fig. 3d, e). Conversely, the genetic determinants of the myeloid response were much more virus-dependent (49% of myeloid reQTLs, interaction *p*-value < 0.01), with 46 and 185 reQTLs displaying specific, stronger effects following stimulation with SARS-CoV-2 and IAV, respectively. The strongest SARS-CoV-2-specific reQTL (rs534191, Student’s *p*-value = 1.96 × 10^-16^ for COV and 0.05 for IAV; Fig. 3f) was identified in myeloid cells, at the *MMP1* locus, which encodes a reported biomarker of COVID-19 severity^37^. These analyses revealed that the genetic bases of leukocyte responses to SARS-CoV-2 are highly cell type-dependent, with the myeloid response being strongly virus-specific.

### Ancestry effects on immune response variation

We then evaluated the contribution of genetic ancestry to population differences in immune responses, by focusing on popDEGs and popDRGs. We found that 11-24% of the genes expressed genome-wide had at least one eQTL, but this proportion increased to up to 56% and 60% for popDEGs and popDRGs, respectively, not explained by cellular heterogeneity (Fig. 3g, Supplementary Fig. 8c). Furthermore, the popDEGs and popDRGs displaying the largest population differences were more likely to be under genetic control and associated with eQTLs/reQTLs with the largest effect sizes (Supplementary Fig. 8d-f). We used mediation analysis to assess, for each gene, cell lineage and virus treatment, the fraction of population differences explained by genetics (i.e., the most significant eQTL) or cellular heterogeneity (Supplementary Table 5). Cellular composition had a broad effect on population differences in gene expression and responses to viral stimuli (explaining 16-62% of population differences per lineage and virus condition, with the strongest effect in NK cells), whereas genetic variants had a weaker overall effect (accounting for 13-35% of population differences; Fig. 3h, Supplementary Fig. 8g). However, genetic variants had strong effects on a subset of genes (141-433 genes per lineage) for which they accounted for 32-58% of population differences in expression. For example, 81-100% of the difference in *GBP7* expression between donors of African and European ancestry were explained by a single variant displaying strong population differentiation (rs1142888, derived allele frequency (DAF) = 0.13 and 0.53 in African- and European-ancestry individuals, respectively, *F*_ST_ = 0.26, |β_eQTL_| > 1.7 across lineages upon stimulation). Thus, variation in immune responses across populations is driven largely by cellular heterogeneity, but common *cis*-genetic variants that present marked allele frequency variation contribute to population differences at specific loci.

### Natural selection and SARS-CoV-2 responses

We explored the contribution of natural selection to population differentiation of immune responses. We first searched for overlaps between eQTLs or reQTLs and genome-wide signals of local adaptation, measured by the population branch statistic (PBS)^38^. We identified 1,616 eQTLs (1,215 genes) and 180 reQTLs (166 genes) displaying a strong allele frequency differentiation (empirical *p*-value < 0.01) in at least one population (Supplementary Table S6a). They included key players in IFN-mediated antiviral immunity, such as *DHX58* and *TRIM14* in African-ancestry individuals, *ISG20, IFIT5, BST2* and *IFITM2-3* in European-ancestry individuals, and *IFI44L* and *IFITM2* in East Asian-ancestry individuals. We then used CLUES^39^ to identify rapid changes in the frequency trajectory of (r)eQTLs over the last 2,000 generations (i.e., 56,000 years) in each population (Supplementary Fig. 9a-d, Supplementary Table S6b). We found signals of rapid adaptation (max. |*Z*| > 3, Methods) targeting the same (*IFITM2, IFIT5*) or different (*ISG20, IFITM3, TRIM14*) eQTLs at highly differentiated genes. We determined whether selection had altered gene expression in specific cell types or in response to SARS-CoV-2 or IAV, by testing for an increase in population differentiation (PBS) at specific eQTLs/reQTLs, relative to random SNPs matched for allele frequency, linkage disequilibrium (LD) and distance to the nearest gene. In the basal state, eQTLs were more strongly differentiated in Europeans, with the strongest signal observed for γδ T cells (Fig. 4a, Supplementary Fig. 9e). We found that 34% of popDEGs — for which genetics was found to mediate > 50% of the differences between donors of African and European ancestries — were associated with signals of rapid adaptation in Europeans (*vs*. 21% in Africans, Fisher’s exact *p*-value = 7.7 × 10^-6^). For example, population differences at *GBP7* have been driven by a rapid frequency increase, over the last 782-1,272 generations, of the rs1142888-G allele in Europeans (max. |*Z*| > 4.3, Supplementary Fig. 9f).

**Figure 4.**
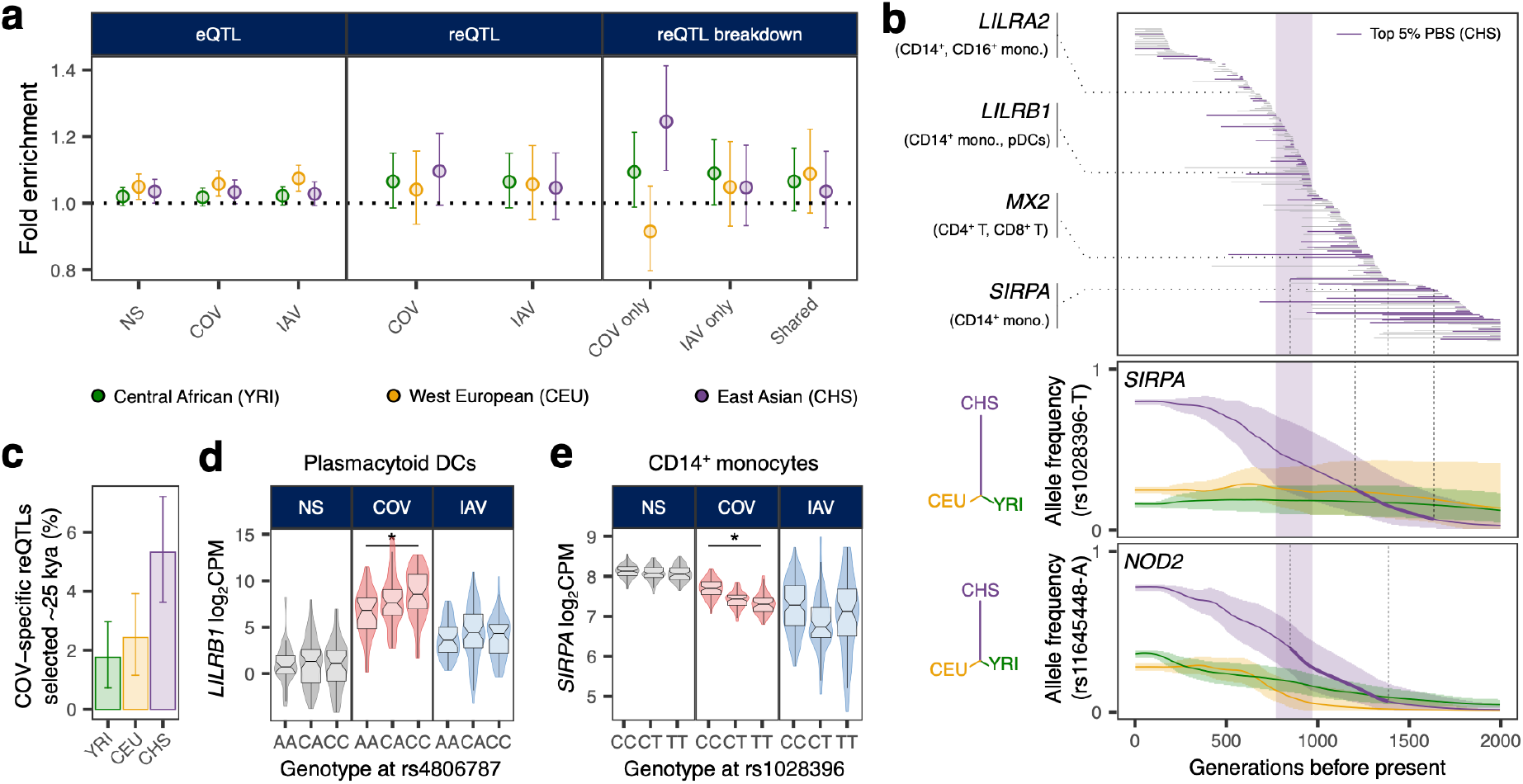
Natural selection effects on population differentiation of immune responses. **a,** Fold-enrichment in local adaptation signals, defined by the population branch statistic (PBS), and 95% CI, for eQTLs and reQTLs relative to randomly selected variants, matched for minor allele frequency, distance to nearest gene, and LD score. **b**. Estimated periods of selection, over the past 2,000 generations, for 245 SARS-CoV-2 reQTLs with significant rapid adaptation signals in East Asians (CHS) (max. |*Z*-score| > 3). Variants are ordered in descending order of time to the onset of selection. The area shaded in purple highlights a period corresponding to 770 to 970 generations ago that has been shown to be associated with polygenic adaptation signals at host coronavirus-interacting proteins in East Asians^23^. Several immunity-related genes are highlighted (top panel). Allele frequency trajectories of two SARS-CoV-2 reQTLs (rs1028396 at *SIRPA* and rs11645448 at *NOD2*) in Central Africans (YRI, green), West Europeans (CEU, yellow) and East Asians (CHS, purple). Shaded areas indicate the 95% CIs. Dendrograms show the estimated population phylogeny for each eQTL based on PBS (i.e., the branch length between each pair of populations is proportional to - log_10_(1-F_ST_;)). **c,** Percentage of SARS-CoV-2-specific reQTLs presenting selection signals in different populations, between 770 and 970 generations ago, with resampling-based 95% CIs. **d, e,** Examples of SARS-CoV-2-induced reQTLs at *LILRB1* (rs4806787) in plasmacytoid dendritic cells and *SIRPA* (rs1028396) in CD14^+^ monocytes. *Student’s *t*-test *p*-value < 0.01; middle line: median; notches: 95% CI of median, box limits: upper and lower quartiles; whiskers: 1.5× interquartile range; points: outliers.

Focusing on the response to viral stimuli, we found that SARS-CoV-2 reQTLs were enriched in signals of population differentiation, specifically in East Asians (fold-enrichment (FE) = 1.24, one-sided resampling *p*-value < 2×10^-4^, Fig. 4a). Furthermore, among SARS-CoV-2-specific reQTLs, 28 reQTLs (5.3%) displayed signals of rapid adaptation (max. |*Z*| > 3) in East Asians starting 770-970 generations ago (~25,000 years) – a time frame associated with polygenic adaptation at SARS-CoV-2-interacting proteins^23^ (OR relative to other populations = 2.6, Fisher’s exact *p*-value = 7.3 × 10^-4^, Fig. 4b, c, Supplementary Fig. 9g, h). A noteworthy example is the immune mediator *LILRB1*, in which we detected a SARS-CoV-2-specific reQTL (rs4806787) in pDCs (Fig. 4d). However, the selection events making the largest contribution to the differentiation of SARS-CoV-2 immune responses in East Asia (top 5% PBS) began before this time period (> 970 generations ago, OR = 1.94, Fisher’s exact *p*-value = 0.019, Fig. 4b). For example, the rs1028396-T allele, associated with a weaker response of *SIRPA* to SARS-CoV-2 in CD14^+^ monocytes (80% in East Asia *vs*. 16-25% elsewhere) is characterized by a selection signal beginning more than 45,000 years ago (Fig. 4b, e). SIRPα has been shown to inhibit infection by endocytic viruses, such as SARS-CoV-2 (ref.^40^). These results are consistent with a history of recurrent genetic adaptation targeting antiviral immunity over the last 50,000 years and contributing to present-day population differences in SARS-CoV-2 immune responses.

### Functional consequences of Neanderthal introgression

We investigated the effect of the introgression of genetic material from archaic humans, such as Neanderthals or Denisovans, on present-day immune responses to viral challenges, by defining a set of 100,345 introgressed ‘archaic’ alleles (aSNPs) and determining whether eQTLs were over/underrepresented among introgressed variants relative to random, matched SNPs (Methods). We found that archaic haplotypes were 1.3-1.4 times more likely to alter gene expression, in the basal state (one-sided permutation *p*-value = 0.02) and after stimulation with SARS-CoV-2 or IAV (one-sided permutation *p*-value = 5×10^-4^ and 6×10^-3^, respectively) in Europeans, whereas this trend was not significant in East Asians (FE = 1.1, one-sided permutation *p*-value > 0.09, for all sets of conditions, Fig. 5a, Supplementary Table S7a-c). Enrichment was strongest in SARS-CoV-2-stimulated CD16^+^ and IAV-infected CD14^+^ monocytes, suggesting that archaic haplotypes altering myeloid responses to viruses have been preferentially retained in the genomes of modern Europeans. Furthermore, archaic haplotypes regulating gene expression are present at higher frequencies than archaic haplotypes without eQTLs in Europeans, after adjustment for the mean of minor allele frequencies worldwide to ensure similar power for the detection of eQTLs (Δ *f*(introgressed allele) > 4.1%, Student’s *p*-value < 1.5×10^-8^; Methods, Fig. 5b, Supplementary Table S7d, e), providing evidence for the adaptive nature of Neanderthal introgression.

**Figure 5.**
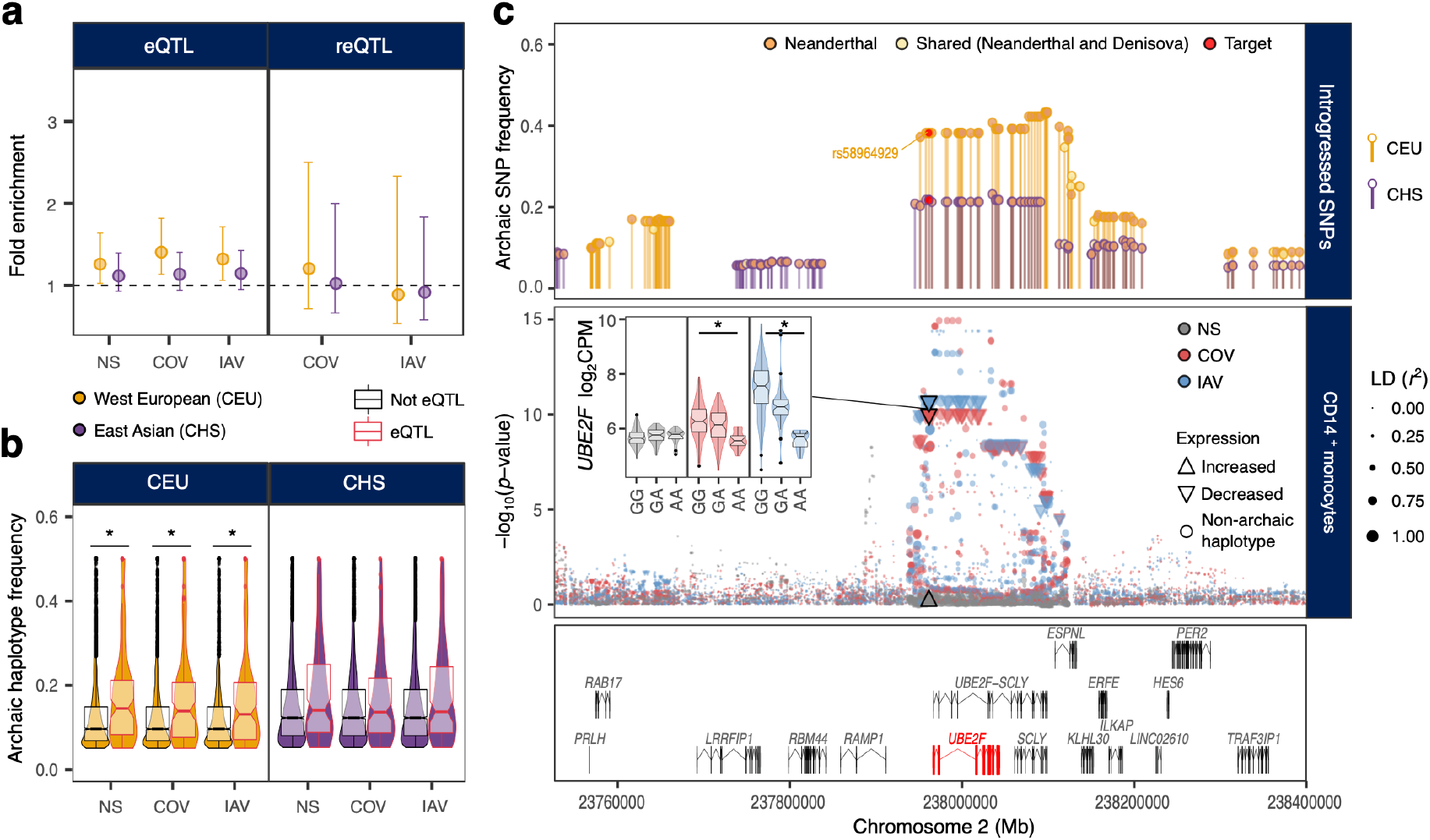
Impact of archaic introgression on molecular and cellular phenotypes. **a**, Enrichment of eQTLs and reQTLs in introgressed haplotypes. For each comparison, the mean observed/expected ratio and the 95% CIs are reported (based on 10,000 resamplings). **b**, For each population and stimulation condition, the frequencies of introgressed haplotypes are compared according to their effects on gene expression (eQTL *vs*. non-eQTL; *Wilcoxon’s *p*-value < 0.001; middle line: median; notches: 95% CI of median, box limits: upper and lower quartiles; whiskers: 1.5× interquartile range; points: outliers). **c**, Example of adaptive introgression at the *UBE2F* locus (reQTL rs58964929). Upper panel: frequency and nature of archaic alleles across the genomic region of chromosome 2 containing *UBE2F*. Each dot represents an allele of archaic origin, and its color indicates whether it was present in the Vindija Neanderthal genome (orange) or was common to the Vindija Neanderthal and Denisova genomes (light yellow). The eQTL index SNP is shown in red. The frequency in West Europeans (CEU, yellow) and East Asians (CHS, purple) is indicated on the *y*-axis. Middle panel: monocyte eQTL *p*-values for SNPs at the *UBE2F* locus, color-coded according to stimulation conditions (gray: non-stimulated (NS), red: SARS-CoV-2-stimulated (COV), blue: IAV-stimulated (IAV)). Each dot represents a SNP and its size (area) is proportional to the LD (*r^2^*) values between the SNP and nearby archaic alleles. For the archaic alleles, arrows indicate the effect of the allele on gene expression. Lower panel: gene structure in the chromosome 2 region, with the *UBE2F* gene highlighted in red.

We characterized the functional consequences of archaic introgression at the level of individual cell types, by focusing on introgressed eQTLs where the archaic allele was found at its highest frequency in Eurasians (i.e., 5% of most frequent SNPs). These eQTLs included known adaptively introgressed variants at *OAS1-3* or *PNMA1* in Europeans and *TLR1, FANCA* or *IL10RA* in East Asians^17,41–44^, for which we delineated the cellular and molecular effects (Supplementary Fig. 10a, b, Supplementary Table S7f). For example, the COVID-19-associated variant rs10774671 at *OAS1* (ref.^45^) exerts its strongest effect in IAV-stimulated γδ T cells and two alleles in the *TLR1-6-10* region (rs189688666-T and rs112318878-T) have opposite effects on *TLR1* expression in IAV-stimulated monocytes and resting CD4^+^ T cells.

We also identified previously unreported signals of Neanderthal introgression affecting immunity phenotypes. For example, an introgressed eQTL (rs11119346-T, 43% in East Asians *vs*. < 3% in Europeans) was found to downregulate *TRAF3IP3 —* encoding a negative regulator of the cytosolic RNA-induced IFN response^46^ — specifically in IAV-infected monocytes, thereby favoring IFN release after viral infection (Supplementary Fig. 10c, d). Likewise, a 35.5-kb Neanderthal haplotype with a frequency of 61% in East Asians (*vs*. 24% in Europeans) contains the rs9520848-C allele, which is associated with higher basal expression for the cytokine gene *TNFSF13B* in MAIT cells (Supplementary Fig. 10e, f). We also identified an introgressed reQTL (rs58964929-A) at *UBE2F* that was present in 38% of Europeans (*vs*. 22% of East Asians) and decreased *UBE2F* responses to SARS-CoV-2 and IAV in monocytes (Fig. 5c). UBE2F is involved in neddylation, a posttranslational modification required for the nuclear translocation of IRF7 by myeloid cells following infection with RNA viruses and, thus, for the induction of type I IFN responses^47^. Collectively, these results document the molecular and cellular mechanisms through which archaic introgression has altered immune functions.

### Immunity-related eQTLs and COVID-19 risk

We investigated the contributions of genetic variants altering responses to SARS-CoV-2 *in vitro* to COVID-19 risk *in vivo*, by determining whether eQTLs/reQTLs were more strongly associated with COVID-19 hits detected by genome-wide association studies^7^ than random, matched SNPs (Methods). We observed an enrichment in eQTLs at loci associated with both susceptibility (reported cases) and severity (hospitalized or critical cases) (FE = 4.1 and FE > 3.8, respectively, one-sided resampling *p*-value < 10^-4^), and a specific enrichment in reQTLs at severity loci (FE > 3.7, one-sided resampling *p*-value < 3 × 10^-3^; Fig. 6a). This trend was observed across most cell lineages (Supplementary Fig. 11a). Colocalization analyses identified 40 genes at which there was a high probability of (r)eQTL colocalization with COVID-19 hits (coloc. PP_H4_ > 0.8; Supplementary Table S8). These included genes encoding direct regulators of innate immunity, such as *IFNAR2* in non-stimulated CD4^+^ and CD8^+^ T cells, *IRF1* in non-stimulated NK and CD8^+^ T cells, *OAS1* in lymphoid cells stimulated with SARS-CoV-2 and IAV, and *OAS3* in SARS-CoV-2-exposed CD16^+^ monocytes (Fig. 6b, c, Supplementary Fig. 11b, c). These results are consistent with a contribution of immunity-related (r)eQTLs to COVID-19 risk.

**Figure 6.**
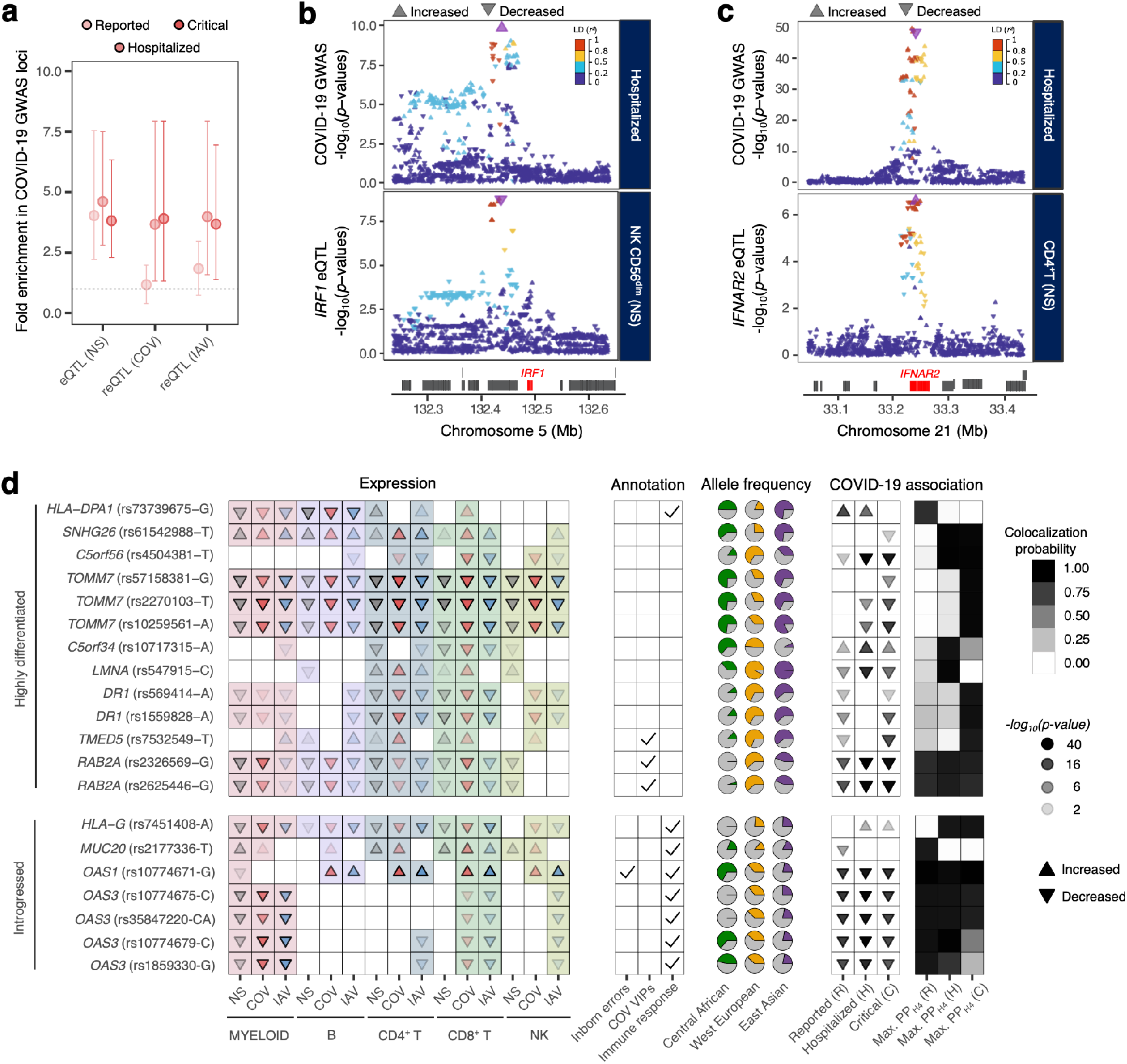
Immunity-related eQTLs and reQTLs contribute to COVID-19 risk. **a**, Enrichment in GWAS loci associated with COVID-19 susceptibility and severity at eQTLs and reQTLs. For each comparison, fold-enrichments and resampling-based 95% confidence intervals are displayed. **b** and **c**, Colocalization of *IRF1* and *IFNAR2* eQTLs with COVID-19 severity loci. Upper panels show the -log_10_(*p*-value) profiles for association with COVID-19-related hospitalization, and lower panels show the -log_10_(*p*-values) profile for association with expression in non-stimulated CD56^dim^ NK cells (*IRF1*) and CD4^+^ T cells (*IFNAR2*). In each panel, the color code reflects the degree of LD (*r*^2^) with the consensus SNP identified in the colocalization analysis (purple). For each SNP, the direction of the arrow indicates the direction of the effect. **d,** Features of (r)eQTLs colocalizing with COVID-19 risk loci (PP_H4_ > 0.8) and presenting either strong population differentiation (top 1% PBS genome-wide) or evidence of Neanderthal introgression. From left to right: (i) effects of the target allele on gene expression across immune lineages and stimulation conditions, (ii) clinical and functional annotations of associated genes, (iii) present-day population frequencies of the target allele, and (iv) effects of the target allele on COVID-19 risk (infection, hospitalization, and critical state) and colocalization probability. Arrows indicate increases/decreases in gene expression or disease risk with each copy of the target allele, and opacity indicates the increases in effect size. In the leftmost panel, arrow colors indicate the stimulation conditions (gray: non-stimulated (NS), red: SARS-CoV-2-stimulated (COV), blue: IAV-stimulated (IAV)). For each eQTL, the target allele is defined as either (i) the derived allele for highly differentiated eQTLs, or (ii) the allele that segregates with the archaic haplotype for introgressed eQTLs in Eurasians. When the ancestral state is unknown, the minor allele is used as a proxy for the derived allele. Note that in some cases (e.g., *OAS1*) the introgressed allele can be present at high frequency in Africa, which is attributed to the reintroduction in Eurasia of an ancient allele by Neanderthals^45^.

Focusing on the evolutionary factors affecting present-day COVID-19 risk, we identified 20 eQTLs that (i) colocalized with COVID-19 susceptibility or severity hits (PP_H4_ > 0.8) and (ii) presented positive selection signals (top 1% PBS, *n* = 13 eQTLs) or evidence of archaic introgression (*n* = 7 eQTLs) (Fig. 6d). For example, we found two variants in high LD (rs569414 and rs1559828, *r*^2^ > 0.73) at the *DR1* locus that displayed extremely high levels of population differentiation that could be attributed to out-of-Africa selection (DAF = 0.13 in Africa *vs*. > 0.62 in Eurasia, Supplementary Fig. 11d). Interestingly, DR1 suppresses type I IFN responses^48^ and the alleles subject to selection, which today decrease COVID-19 severity, reduce *DR1* expression in most immune cells (Fig. 6d). Furthermore, we identified a ~39-kb Neanderthal haplotype spanning the *MUC20* locus in Europeans and East Asians, in which the rs2177336-T allele is associated with both higher levels of *MUC20* expression in SARS-CoV-2-stimulated cells, particularly for CD4^+^ T cells, and a lower susceptibility to COVID-19. Together, these results reveal the contribution of past selection or Neanderthal introgression affecting immune response variation to current disparities in COVID-19 risk.

## Discussion

The degree and sources of the variation of immune responses to SARS-CoV-2 have emerged as major issues since the beginning of the COVID-19 pandemic^4,6,14,15^. Based on single-cell approaches, this study provides evidence that cellular proportions, the variation of which is largely due to environmental exposures, are major drivers of population differences in SARS-CoV-2 immune responses. The higher proportions of memory cells detected in lymphoid lineages in individuals of African descent and their association with persistent CMV infection suggest that population differences in cellular activation states may be driven primarily by lifelong pathogen exposure. This highlights how socio-environmental factors (here, pathogen exposure) may covary with individual ancestry (i.e., genetic background), which may lead to an overestimation of the effects of ancestry on phenotypic variation (i.e., immune responses to SARS-CoV-2). Still, common genetic variants can also contribute to the observed variability of immune responses to viral challenges, but their effects tend to be limited to a subset of genes displaying strong population differentiation. This is best illustrated by the rs1142888-G variant, which solely accounts for the > 2.8-fold higher levels of *GBP7* expression in response to viral stimulation in Europeans than in Africans. The higher frequency of this variant in Europe appears to result from a selection event that occurred 21,900-35,600 years ago. GBP7 has been shown to facilitate IAV replication by suppressing innate immunity^49^, but it also regulates IFN-γ-induced oxidative host defense and confers resistance to intracellular bacteria, such as *Listeria monocytogenes* and *Mycobacterium tuberculosis*^50^, providing a plausible mechanism for the occurrence of positive selection at this locus.

This study also provides evidence to suggest that past natural selection and admixture with Neanderthals contributed to the differentiation of immune responses to SARS-CoV-2. We found traces of a selection event targeting SARS-CoV-2-specific reQTLs ~25,000 years ago in the ancestors of East Asians, coinciding with the proposed timing of an ancient epidemic affecting the evolution of host coronavirus-interacting proteins^23,24^. However, we found little overlap between the alleles selected during this period in East Asia and the reported genetic variants underlying COVID-19 risk, suggesting that there have been changes in the genetic basis of infectious diseases over time, possibly due to the evolution of viruses themselves. Nevertheless, we identified cases (e.g., *DR1, OAS1-3, TOMM7, MUC20*) in which selection or archaic introgression contributed to changes in both immune responses to SARS-CoV-2 and the outcome of COVID-19. Genomic studies based on ancestry-aware polygenic risk scores derived from cross-population GWAS will be required to establish a causal link between past adaptation and present-day population differences in COVID-19.

Finally, dissection of the genetic architecture of immune response variation across a wide range of cell types provides mechanistic insight into the effect of alleles previously associated with COVID-19 risk. We found that several variants of *IRF1, IFNAR2*, and *DR1* associated with lower COVID-19 severity increase type I IFN signaling in lymphoid cells by upregulating *IRF1* and *IFNAR2* or downregulating *DR1*, providing evidence for the importance of efficient IFN signaling for a favorable clinical outcome of SARS-CoV-2 infection^4,11–13^. Another relevant example is provided by the *MUC20* locus, at which we identified a Neanderthal-introgressed eQTL that both increased *MUC20* expression in SARS-CoV-2-stimulated CD4^+^ T cells and has been shown to decrease COVID-19 susceptibility. Given the role of mucins in forming a barrier against infection in the nasal epithelium, we suggest that the Neanderthal haplotype confers greater resistance to viral infections via a similar effect in nasal epithelial cells.

Overall, these findings highlight the value of using single-cell approaches to capture the full diversity of the human immune response to RNA viruses, and SARS-CoV-2 in particular, and shed light on the environmental, genetic and evolutionary drivers of immune response variation across individuals and populations.

## Methods

### Sample collection

The individuals of self-reported African (AFB) and European (EUB) descent studied are part of the EVOIMMUNOPOP cohort^17^. Briefly, 390 healthy male donors were recruited between 2012 and 2013 in Ghent (Belgium), thus before the COVID-19 pandemic (188 of self-reported African descent, and 202 of self-reported European descent). Blood was obtained from the healthy volunteers, and the peripheral blood mononuclear cell (PBMC) fraction was isolated and frozen. Inclusion in the current study was restricted to 80 nominally healthy individuals of each ancestry, between 19 and 50 years of age at the time of sample collection. Donors of African descent originated from West Central Africa, with >90% being born in either Cameroon or the Democratic Republic of Congo. For this study, an additional 71 individuals of East Asian descent (ASH) were included (62 donors left after quality control, see “Single-cell RNA sequencing library preparation and data processing”). ASH individuals were recruited at the School of Public Health, University of Hong Kong, and were included in a community-based sero-epidemiological COVID-19 study (research protocol number JTW 2020.02). Inclusion for the study described here was restricted to nominally healthy ASH individuals (30 men and 41 women) aged between 19 and 65 years of age and seronegative for SARS-CoV-2. Samples were collected at the Red Cross Blood Transfusion Service (Hong Kong) where the PBMC fraction was isolated and frozen.

In this study, we refer to individuals of ‘Central African’ (AFB), ‘West European’ (EUB) and ‘East Asian’ (ASH) ancestries to describe individuals who are genetically similar (i.e., lowest *F*ST values) to populations from West-Central Africa, Western Europe and East Asia, using the 1,000 Genomes (1KG) Project^51^ data as a reference (Supplementary Fig. 1a). Of note, the AFB, EUB and ASH samples present no detectable evidence of recent genetic admixture with populations originating from another continent (e.g., AFB present no traces of recent admixture with EUB).

All samples were collected after written informed consent had been obtained from the donors, and the study was approved by the ethics committee of Ghent University (Belgium, no. B670201214647), the Institutional Review Board of the University of Hong-Kong (no. UW 20-132), and the relevant French authorities (CPP, CCITRS and CNIL). This study was also monitored by the Ethics Board of Institut Pasteur (EVOIMMUNOPOP-281297).

### Genome-wide DNA genotyping

The AFB and EUB individuals were previously genotyped at 4,301,332 SNPs, with the Omni5 Quad BeadChip (Illumina, California) with processing as previously described^17^. The additional 71 ASH donors were genotyped separately at 4,327,108 SNPs with the Infinium Omni5-4 v1.2 BeadChip (Illumina, California). We updated SNP identifiers based on Illumina annotation files (https://support.illumina.com/content/dam/illumina-support/documents/downloads/productfiles/humanomni5-4/v1-2/infinium-omni5-4-v1-2-a1-b144-rsids.zip) and called the genotypes of all ASH individuals jointly on GenomeStudio (https://www.illumina.com/techniques/microarrays/array-data-analysis-experimental-design/genomestudio.html). We then removed SNPs with (i) no “rs” identifiers or with no assigned chromosome or genomic position (*n* = 14,637); (ii) duplicated identifiers (*n* = 5,059); or (iii) a call rate < 95% (*n* = 10,622). We then used the 1KG Project Phase 3 data^51^ as a reference for merging the ASH genotyping data with that of AFB and EUB individuals and detecting SNPs misaligned between the three genotype datasets. Before merging, we removed SNPs that (i) were absent from either the Omni5 or 1KG datasets (*n* = 469,535); (ii) were transversions (*n* = 138,410); (iii) had incompatible alleles between datasets, before and after allele flipping (*n* = 1,250); and (iv) had allele frequency differences of more than 20% between the AFB and Luhya from Webuye, Kenya (LWK) and Yoruba from Ibadan, Nigeria (YRI), or between the EUB and Utah residents with Northern and Western European ancestry (CEU) and British individuals from England and Scotland (GBR), or between the ASH and Southern Han Chinese (CHS) (*n* = 777). Once the data had been merged, we performed principal components analysis (PCA) with *PLINK 1.9* (ref.^52^) and ensured that the three study populations (i.e., AFB, EUB, and ASH) overlapped with the corresponding 1KG populations, to exclude batch effects between genotyping platforms (Supplementary Fig. 1a). The final genotyping dataset included 3,723,840 SNPs.

### Haplotype phasing and imputation

After merging genotypes from AFB, EUB and ASH donors, we filtered genotypes for duplicates with *bcftools norm --rm-dup all* (v1.16) (ref.^53^) and lifted all genotypes over to the human genome assembly GRCh38 with GATK’s (v4.1.2.0) *LiftoverVcf* using the *RECOVER_SWAPPED_ALT_REF=TRUE* option^54^. We then filtered out duplicated variants again before phasing genotypes with *SHAPEIT4* (v4.2.1) (ref.^55^) and imputing missing variants with *Beagle5.1* (version: 18May20.d20) (ref.^56^), treating each chromosome separately. For both phasing and imputation, we used the genotypes of 2504 unrelated individuals from the 1,000 Genomes (1KG) Project Phase 3 data as a reference (downloaded from ftp://ftp/1000genomes.ebi.ac.uk/vol1/ftp/release20130502 and lifted over to GRCh38) and downloaded genetic maps from the GitHub pages of the associated software (i.e., *SHAPEI4* for phasing and *Beagle5.1* for imputation). A third round of duplicate filtering was performed after phasing and before imputation with *Beagle5.1* (version: 18May20.d20) (ref.^56^). Phasing was performed setting *-pbwt-depth=8* and imputation was performed assuming an effective population size (*N*_e_) of 20,000. The quality of imputation was assessed by cross-validation; specifically, we performed 100 independent rounds of imputation excluding 1% of the variants and compared the imputed allelic dosage with the observed genotypes for these variants (Supplementary Fig. 1b, c). The results obtained confirmed that imputation quality was satisfactory, with 98% of common variants (i.e., MAF > 5%) having an *r*^2^ > 0.8 for the correlation between observed and imputed genotypes (>95% concordance for 96% of common variants). Following imputation, variants with a MAF < 1% or with a low predicted quality of imputation (i.e., DR2 < 0.9) were excluded, yielding a final dataset of 13,691,029 SNPs for downstream analyses.

### Viruses used in this study

The SARS-CoV-2 reference strain used in this study (BetaCoV/France/GE1973/2020) was supplied by the National Reference Centre for Respiratory Viruses hosted by Institut Pasteur (Paris, France) and headed by Dr. Sylvie van der Werf. The human sample from which the strain was isolated was provided by Dr. Laurent Andreoletti from the Robert Debré Hospital (Paris, France). The influenza A virus strain used in this study (IAV, PR/8, H1N1/1934) was purchased from Charles River laboratories (lot n° #3X051116) and provided in ready-to-use aliquots that were stored at −80°C.

### SARS-CoV-2 stock production

To produce SARS-CoV-2, we used African green monkey kidney Vero E6 cells that were maintained at 37°C in 5% CO_2_ in Dulbecco’s minimum essential medium (DMEM) (Sigma-Aldrich) supplemented with 10% fetal bovine serum (FBS, Dutscher) and 1% penicillin/streptomycin (P/S, Gibco, Thermo Fisher Scientific). Vero E6 cells were plated at 80% confluence in 150 cm^2^ flasks and infected with SARS-CoV-2 at a multiplicity of infection (MOI) of 0.01 in DMEM supplemented with 2% FBS and 1% P/S. After 1 hour, the inoculum was removed and replaced with DMEM supplemented with 10% FBS, 1% P/S, and cells were incubated for 72 hours at 37°C in 5% CO_2_. The cell culture supernatant was collected and centrifuged for 10 min at 3,000 r.p.m to remove cellular debris, and polyethylene glycol (PEG; PEG8000, Sigma-Aldrich) precipitation was performed to concentrate the viral suspension. Briefly, 1 L of viral stock was incubated with 250 mL of 40% PEG solution (i.e., 8% PEG final) overnight at 4°C. The suspension was centrifuged at 10,000*g* for 30 minutes at 4°C and the resulting pellet was resuspended in 100 mL of RPMI medium (Gibco, Thermo Fisher Scientific) supplemented with 10% FBS (referred to as R10) and viral aliquots were stored at −80°C. SARS-CoV-2 viral titers were determined by focus-forming unit (FFU) assay as previously described^57^. Briefly, Vero-E6 cells were plated in a 96-multiwell plate with 2 × 10^4^ cells per well. The cellular monolayer was infected with serial dilutions (1:10) of viral stock and overlaid with a semi-solid 1.5% carboxymethylcellulose (CMC, Sigma-Aldrich) and 1x MEM medium for 36 hours at 37°C. Cells were then fixed with 4% paraformaldehyde (Sigma-Aldrich), and permeabilized with 1× PBS–0.5% Triton X-100 (Sigma-Aldrich). Infectious foci were stained with a human anti-SARS-CoV-2 Spike antibody (H2-162, Hugo Mouquet’s laboratory, Institut Pasteur) and the corresponding HRP-conjugated secondary antibody (Sigma-Aldrich). Foci were visualized by 3,3’-diaminobenzidine staining solution (DAB, Sigma-Aldrich) staining and counted with the BioSpot suite of a C.T.L. ImmunoSpot S6 Image Analyzer.

### *In vitro* peripheral blood mononuclear cell stimulation

We performed single-cell RNA-sequencing on SARS-CoV-2-, IAV- and mock-stimulated (referred to as “non-stimulated” condition) PBMCs from healthy donors (80 AFB, 80 EUB and 71 ASH) in 16 experimental runs. We first performed a kinetic experiment (run 1) on samples from 4 AFB and 4 EUB stimulated for 0, 6 and 24 hours to validate our *in vitro* model across different time points (Supplementary Fig. 2, Supplementary Table 1). The 6-hour time point was identified as the optimal time point for the analysis (Supplementary Note 1). We then processed the rest of the cohort, over runs 2 through 15. Finally, we reprocessed some samples (run 16) to assess technical variability in our setting and to increase *in silico* cell counts (see ‘ Single-cell RNA sequencing library preparation and data processing’ section). Ancestry-related batch effects were minimized by scheduling sample processing to ensure a balanced distribution of AFB, EUB and ASH donors within each run.

For each run, cryopreserved PBMCs were thawed in a 37°C water bath, transferred to 25 mL of R10 medium (i.e., RPMI 1640 supplemented with 10% heat-inactivated fetal bovine serum) at 37°C, and centrifuged at 300*g* for 10 minutes at room temperature. Cells were counted, re-suspended at 2× 10^6^ cells/mL in warm R10 in 25cm^2^ flasks, and rested overnight (i.e., 14 hours) at 37°C. The next morning, PBMCs were washed and re-suspended at a density of 3.3×10^6^ cells/mL in R10; 120 μL of a suspension containing 4×10^5^ cells from each sample was then plated in a 96-well untreated plate (Greiner Bio-One) for each of the three sets of stimulation conditions. We added 80 μL of either R10 (non-stimulated), SARS-CoV-2 or IAV stock (corresponding to 4×10^5^ focus-forming units diluted in R10) to the cells, so as to achieve a multiplicity of infection (MOI) of 1 and an optimal PBMC concentration of 2× 106 cells/mL. Cells were incubated at 37°C for 0, 6 or 24 hours for the kinetic experiment (run 1), and for 6 hours for all subsequent runs (runs 2 to 16), in a biosafety level 3 (BSL-3) facility at Institut Pasteur, Paris. The plates were then centrifuged at 300*g* for 10 minutes and supernatants were stored at −20°C until use (see ‘Supernatant cytokine assays’ section). All samples from the same run were resuspended in Dulbecco’s phosphate-buffered saline (PBS, Gibco), supplemented with 0.04% bovine serum albumin (BSA, Miltenyi Biotec), and multiplexed in eight pools according to a pre-established study design (Supplementary Fig. 3a, Supplementary Table 2a). The cells from each pool were counted with a Cell Countess II automated cell counter (Thermo Fisher Scientific) and cell density was adjusted to 1,000 viable cells/μL 0.04% BSA in PBS.

### Single-cell RNA sequencing library preparation and data processing

We generated scRNA-seq cDNA libraries with a Chromium Controller (10X Genomics) according to the manufacturer’s instructions for the Chromium Single Cell 3’ Library and Gel Bead Kits (v3.1). Library quality and concentration were assessed with an Agilent 2100 Bioanalyzer and a Qubit fluorometer (Thermo Fisher Scientific). The final products were processed for high-throughput sequencing on a HiSeqX platform (Illumina Inc.).

Paired-end sequencing reads from each of the 133 scRNA-seq cDNA libraries (13 libraries from the kinetic experiment and 120 from the population-level study) were independently mapped onto the concatenated human (GRCh38), SARS-CoV-2 (hCoV-19/France/GE1973/2020) and IAV (A/Puerto Rico/8/1934(H1N1)) genome sequences with the *STARsolo* aligner (v2.7.8a) (ref.^58^) (Supplementary Fig. 3b). We obtained a mean of 10,785 cell-containing droplets per library, and each droplet was assigned to its sample of origin with *Demuxlet* (v0.1) (ref.^59^), based on the genotyping data available for each individual. Singlet/doublet calls were compared with the output of *Freemuxlet* (v0.1) (ref.^59^) to ensure good agreement (Supplementary Fig. 3c-e). We loaded feature-barcode matrices for all cell-containing droplets identified as singlets by *Demuxlet* in each scRNA-seq library onto a *SingleCellExperiment* (v1.14.1) object^60^. Data from barcodes associated with low-quality or dying cells were removed with a hard threshold-based filtering strategy based on three metrics: cells with fewer than 1,500 total UMI counts, 500 detected features or a mitochondrial gene content exceeding 20% were removed from each sequencing library (Supplementary Fig. 3f). We also discarded samples nine ASH donors from whom fewer than 500 cells were obtained in at least one condition (Supplementary Fig. 3g).

We then log-normalized raw UMI counts with a unit pseudocount and library size factors (i.e., number of reads associated with each barcode) were calculated with *quickClusters* and *computeSumFactors* from the *scran* package (v1.20.1) (ref.^60^). We then calculated the mean and variance of log counts for each gene and broke the variance down into a biological and a technical component with the *fitTrendPoisson* and *modelGeneVarByPoisson* functions of *scran*. This approach assumes that technical noise is Poisson-distributed and simulates Poisson-distributed data to derive the mean-variance relationship expected in the absence of biological variation. Excess variance relative to the null hypothesis is considered to correspond to the biological variance. We retained only those genes for which the biological variance component was positive with a FDR below 1%. We used this filtered feature set and the technical variance component modeled from the data to run principal components analysis (PCA) with *denoisePCA* from *scran*, thus discarding later components more likely to capture technical noise. Doublets (i.e., barcodes assigned to cells from different individuals captured in the same droplet) are likely to be in close neighborhoods when projected onto a subspace of the data of lower dimensionality (Supplementary Fig. 3h). We therefore used a *k*-nearest neighbors (*k*-NN) approach to discard cryptic doublets (i.e., barcodes associated to different cells from the same individual captured in the same droplet). Barcodes identified as singlets by *Demuxlet* but having over 5/25 doublet NNs in the PCA space, were re-assigned as doublets and excluded from further analyses.

Following data preprocessing, we performed a second round of UMI count normalization, feature selection and dimensionality reduction on the cleaned data, to prevent bias due to the presence of low-quality cells and cryptic doublets. Sequence depth differences were equalized between batches (i.e., sequencing libraries) with *multiBatchNorm* from *batchelor* (v1.8.1) to scale library size factors according to the ratio of mean counts between batches^61^ (Supplementary Fig. 3i). We accounted for the different mean-variance trends in each batch, by applying *modelGeneVarByPoisson* separately for each sequencing library, and then combining the results for all batches with *combineVar* from *scran* (ref.^60^). We then bound all 133 separate preprocessed feature-barcode matrices into a single merged *SingleCellExperiment* object, log-normalized UMI counts according to the scaled size factors and selected genes with mean log-expression values over 0.01 or a biological variance compartment exceeding 0.001 (Supplementary Fig. 3j). Based on this set of highly variable genes and the variance decomposition, we then performed PCA on the whole data set with *denoisePCA*, and then used *Harmony* (v0.1.0) on the PCs to adjust for library effects^62^.

### Clustering and cell-type assignment

We performed cluster-based cell-type identification in each stimulation condition, according to a four-step procedure. We first performed low-resolution (*res. parameter*=0.8) shared nearest-neighbors graph-based (k=25) clustering with *FindClusters* from *Seurat* (v4.1.1) with assignment to one of three meta-clusters (i.e., myeloid, B lymphoid and T/NK lymphoid) based on the transcriptional profiles of the cells for canonical markers (e.g., *CD3E-F, CD14, FCGR3A, MS4A1*) (Supplementary Fig. 4a, b). We then performed a second round of clustering at higher resolution (*res. parameter=3*) within each meta-cluster and stimulation condition (Supplementary Fig. 4c). We systematically tested for differential expression between each cluster and the other clusters of the same meta-cluster and stimulation condition. This made it possible to define unbiased markers (|log_2_FC| ≠ 0, FDR < 0.01) for each cluster (Supplementary Fig. 4d). We then used these expression profiles of these genes to assign each cluster manually to one of 22 different *cell types* (Supplementary Fig. 4e), which, for some analyses, were collapsed into five major immune *lineages*. This step was performed in parallel by three investigators to consolidate consensus assignments. We also used cellular indexing of transcriptomes and epitopes by sequencing (CITE-seq) data, generated for a subset of cells (2% of the whole data set), to validate our assignments and redefine clusters presenting ambiguous transcriptional profiles (e.g., memory-like NK cells, Supplementary Fig. 4f).

By calling cell types from high-resolution, homogeneous clusters, assigned independently for each lineage and stimulation condition (i.e., non-stimulated, SARS-CoV-2, and IAV), we were able to preserve much of the diversity in our data set, while avoiding potential confounding effects due to the stimulation conditions. However, some clusters were characterized by markers associated with different cell types. Most of these clusters corresponded to mixtures of similar cell types (e.g., the expression of *CD3E, CD8A, NKG7, CD16* suggested a mixture of cytotoxic CD8^+^ T and NK cells) and were consistent with the known cell hierarchy. Other, less frequent clusters expressed a combination of markers usually associated with lineages originating from different progenitors (e.g., *CD3E* and *CD19*, associated with T and B lymphocytes, respectively). These clusters were considered incoherent and discarded. In the fourth and final step of our procedure, we used linear discriminant analysis to resolve the mixtures that were consistent with the established cell hierarchy, to obtain a final cell assignment (Supplementary Fig. 4g, h). For clusters of mixed identity AB, we built a training data set from 10,000 observations sampled from the set of cells called as A or B, preserving the corresponding frequencies of these cells in the whole dataset. We then used a model trained on these data to predict the specific cellular identities within the mixed cluster.

### Cellular indexing of transcriptomes and epitopes by sequencing

As a means to confirm the identity of specific cell types expressing ambiguous markers at the RNA level, during the last experimental run (run 16), half the cells from each experimental condition were used to perform CITE-seq, according to the manufacturer’s instructions (10X Genomics). PBMCs were washed, resuspended in chilled 1% BSA in PBS and incubated with human TruStain FcX blocking solution (BioLegend) for 10 minutes at 4°C. Cells were then stained with a cocktail of TotalSeq^TM^-B antibodies (BioLegend) previously centrifuged at 14,000*g* for 10 minutes (Supplementary Table S2b). The cells were incubated for 30 minutes at 4°C in the dark and were then washed three times. Cell density was then adjusted to 1,000 viable cells/μL in 1% BSA in PBS. We generated scRNA-seq libraries and cell protein libraries (L131-L134) with the Chromium Single Cell 3’ Reagent Kit (v3.1), using the Feature Barcoding technology for Cell Surface Proteins (10X Genomics).

### Supernatant cytokine assays

Before protein analysis, sample supernatants were treated in the BSL-3 facility to inactivate the viruses, according to a published protocol for SARS-CoV (ref.^63^), which we validated for SARS-CoV-2. Briefly, all samples were treated with 1% (v/v) TRITON X100 (Sigma-Aldrich) for 2 hours at room temperature, which effectively inactivated both SARS-CoV-2 and IAV. Protein concentration was then determined with a commercial Luminex multianalyte assay (Biotechne, R&D Systems) and the SIMOA Homebrew assay (Quanterix). For the Luminex assay, we used the XL Performance Kit according to the manufacturer’s instructions, and proteins were determined with a Bioplex 200 (Bio-Rad). Furthermore, IFN-α, IFN-*γ* (duplex) and IFN-β (single-plex) protein concentrations were quantified in SIMOA digital ELISA tests developed as Quanterix Homebrews according to the manufacturer’s instructions (https://portal.quanterix.com/). The SIMOA IFN-α assay was developed with two autoantibodies specific for IFN-α isolated and cloned (Evitria, Switzerland) from two APS1/APECED patients^64^ and covered by patent application WO2013/098419. These antibodies can be used for the quantification of all IFN-α subtypes with a similar sensitivity. The 8H1 antibody clone was used to coat paramagnetic beads at a concentration of 0.3 mg/mL for use as a capture antibody. The 12H5 antibody was biotinylated (biotin/antibody ratio = 30:1) and used as the detector antibody, at a concentration of 0.3 μg/mL. The SBG enzyme for detection was used at a concentration of 150 pM. Recombinant IFNα17/αI (PBL Assay Science) was used as calibrator. For the IFN-*γ* assay, the MD-1 antibody clone (BioLegend) was used to coat paramagnetic beads at a concentration of 0.3 mg/mL for use as a capture antibody. The MAB285 antibody clone (R&D Systems) was biotinylated (biotin/antibody ratio = 40:1) and used as the detector antibody at a concentration of 0.3 μg/mL. The SBG enzyme used for detection was used at a concentration of 150 pM. Recombinant IFN-*γ* protein (PBL Assay Science) was used as a calibrator. For the IFN-β assay, the 710322-9 IgG1, kappa, mouse monoclonal antibody (PBL Assay Science) was used to coat paramagnetic beads at a concentration of 0.3 mg/mL, for use as a capture antibody. The 710323-9 IgG1 kappa mouse monoclonal antibody was biotinylated (biotin/antibody ratio = 40:1) and used as the detector antibody at a concentration of 1 μg/mL. The SBG enzyme for detection was used at a concentration of 50 pM. Recombinant IFN-β protein (PBL Assay Science) was used as a calibrator. The limit of detection (LOD) of these assays was 0.8 fg/mL for IFN-α, 20 fg/mL for IFN-*γ*, and 0.2 pg/mL for IFN-β, considering the dilution factor of 10.

### Flow cytometry

Frozen PBMCs from three AFB (CMV^+^) and six EUB (three CMV^+^, three CMV^-^) donors were thawed and allowed to rest overnight, as previously described. For each donor, 10^6^ cells were resuspended in PBS supplemented with 2% fetal bovine serum and incubated with human Fc blocking solution (BD Biosciences) for 10 minutes at 4°C. Cells were then stained with the following antibodies for 30 minutes at 4°C: CD3 VioGreen (clone BW264/56, Miltenyi Biotec), CD14 V500 (clone M5E2, BD Biosciences), CD57 Pacific Blue (clone HNK-1, Biolegend), NKp46 PE (clone 9E2/NKp46, BD Biosciences), CD16 PerCP-Cy5.5 (clone 3G8, BD Biosciences), CD56 APC-Vio770 (clone REA196, Miltenyi Biotec), NKG2A FITC (clone REA110, Miltenyi Biotec), NKG2C APC (clone REA205, Miltenyi Biotec). The cells were then washed and acquired on a MACSQuant cytometer (Miltenyi Biotec), and the data were analyzed with *FlowJo* software (v10.7.1) (ref.^65^).

### Cytomegalovirus IgG ELISA

We determined the cytomegalovirus (CMV) serostatus of AFB (*n* = 80) and EUB (*n* = 80) donors with a human anti-IgG CMV ELISA kit (Abcam) on plasma samples, according to the manufacturer’s instructions.

### Quantification of batch effects and replicability

Once all the samples had been processed, we used the kBET metric (v0.99.6) (ref.^66^) to assess the intensity of batch effects and to quantify the relative effects of technical and biological variation on cell clustering. This made it possible to confirm that the variation across libraries, and across experimental runs, remained limited relative to the variation across individuals or across conditions (Supplementary Fig. 5a). We used technical replicates to assess the replicability of our observations across independent stimulations. Agreement was good between the cell proportions and the interferon-stimulated gene (ISG) activity scores inferred across independent runs (r > 0.82, *p* < 7.6 × 10^-13^) (Supplementary Fig. 5b, c).

### Pseudobulk estimation, normalization, and batch correction

Individual variation in gene expression was quantified at two resolutions: five major immune *lineages* and 22 *cell types*. We aggregated raw UMI counts from all high-quality single-cell transcriptomes (*n* = 1,047,824) into bulk expression estimates by summing gene expression values across all cells assigned to the same lineage/cell type and sample (i.e., individual and stimulation conditions) using the *aggregateAcrossCells* function of *scuttle* (v1.2.1) (ref.^67^). We then normalized the raw aggregated UMI counts by library size, generating 3,330 lineage-wise (222 donors × 3 sets of conditions × 5 lineages) and 14,652 cell type-wise (666 samples × 22 cell types) pseudobulk counts-per-million (CPM) values, for all genes in our data set. CPM values were then log_2_-transformed, with an offset of 1 to prevent non-finite values and to stabilize variation for weakly expressed genes. Genes with a mean CPM < 1 across all conditions and lineages/cell types were considered to be non-expressed and were discarded from further analyses, leading to a final set of 12,667 genes at the lineage level (12,672 genes when increasing granularity to 22 cell types), including 12 viral transcripts. To quantify the experimental variation induced by experimental run, library preparation and sequencing, and remove unwanted batch effects, we first used the *lmer* function of the *lme4* package (v1.1-27.1) (ref.^68^) to fit a linear model of the following form in each stimulation condition and for each lineage/cell type:

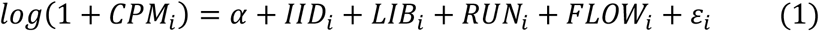

where *CPM_i_* is the gene expression in sample *i* (i.e., one replicate of a given individual and set of experimental conditions), α is the intercept, 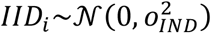 captures the effect of the corresponding individual on gene expression, 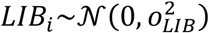 captures the effect of 10X Genomics library preparation, 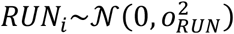 captures the effect of the experimental run, 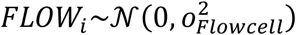 captures the effect of the sequencing flow cell, and *ε_i_* captures residual variation between samples. We then subtracted the estimated value of the library, experimental run, and flow cell effects (as provided by the *ranef* function) from the transformed CPMs of each sample, to obtain batch-corrected CPM values. Finally, we averaged the batch-corrected CPM values obtained across different replicates for the same individual and set of stimulation conditions, to obtain final estimates of gene expression.

For each cell type and stimulation condition, an inverse-normal rank-transformation was applied to the log_2_ CPM of each gene, before testing for differences in gene expression between populations and mapping expression quantitative trait loci. Within each lineage and set of stimulation conditions, we ranked, for each gene, the pseudobulk expression values of all individuals, assigning ranks at random for ties, and replaced each observation with the corresponding quantile from a normal distribution with the same mean and standard deviation as the original expression data. This inverse-normal rank-transformation rendered downstream analyses robust to zero-inflation in the data and outlier values, while maintaining the rank-transformed values on the same scale as the original data.

### Interferon-stimulated gene activity calculation

Interferon-stimulated genes (ISGs) strongly respond to both viruses across all lineages/cell types. We therefore evaluated each donor’s ISG expression level at basal state or upon stimulation with either SARS-CoV-2 or IAV by constructing an “ISG activity” score. For the human genes in our filtered gene set (*N* = 12,655), we defined as ISGs (*n* = 174) those genes included in the union of GSEA’s hallmark (https://www.gsea-msigdb.org/gsea/msigdb/genesets.jsp?collection=H) “IFN-α response” and “IFN-γ response” gene sets, but excluded those from the “inflammatory response” set. We then used *AddModuleScore* from *Seurat* (v4.1.1) (ref.^69^) to measure ISG activity as the mean pseudobulk expression level of ISGs in each sample minus the mean expression for a hundred randomly selected non-ISGs matched for mean magnitude of expression. In all analyses, ISG activity scores were adjusted for cell mortality of the sample by fitting a model of the form:

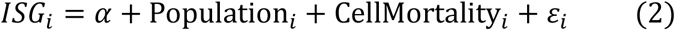

and subtracting the effect of cell mortality from the raw ISG scores. In this model, *LSG_i_* denotes the ISG activity score of individual *i, α* is the intercept, Population_*i*_ and CellMortality_*i*_ are variables capturing the effect of the population, and cell mortality on ISG activity and *ε_i_* are normally distributed residuals. For comparisons with SIMOA-estimated IFN levels, the *carscore* function from the *care* R package^70^ was used to model ISG activity as a function of levels of IFN-α, β, and *γ*, adjusting for population, age, sex and cell mortality. The percentage of ISG variance attributable to each IFN (α, β, or *γ*) was estimated as the square of the resulting CAR scores.

### Modelling population effects on the variation of gene expression

To estimate population effects on gene expression while mitigating any potential batch effect relating to sample processing, we first focused exclusively on AFB and EUB individuals, as all these individuals were recruited during the same sampling campaign and their PBMCs were processed at the same time, with the same experimental procedure^17^. For each immune lineage, cell type, stimulation condition, and gene, we then built a separate linear model of the form:

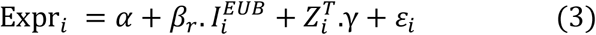

where Expr_*i*_ is the rank-transformed gene expression (log-normalized CPM) for individual *i* in the lineage/cell type and condition under consideration, 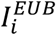 is an indicator variable equal to 1 for European-ancestry individuals and 0 otherwise, and *Z_i_* represents the set of core covariates of the sample that includes the individual’s age and cellular mortality (i.e., proportion of dying cells in each thawed vial, as a proxy of sample quality). In addition, *ε_i_* are the normally distributed residuals and *α, β_r_,γ* are the fitted parameters of the models. In particular, *α* is the intercept, *β_r_* indicates the log_2_fold change difference in expression between individuals of European and African ancestry, and *γ* captures the effects of the set of core covariates on gene expression.

We reasoned that differences in the variance of gene expression between populations might inflate the number of false positives. We therefore used the *vcovHC* function of *sandwich* (v2.5-1) (ref.^71^) with the *type*= *HC3*’ option to compute sandwich estimators of variance that are robust to residual heteroskedasticity. We estimated the *β_r_* coefficients and their standard error with the *coeftest* function of *lmtest* (v0.9-40) (ref. ^72^). FDR was calculated across all conditions and lineages with the Benjamini-Hochberg procedure (*p.adjust* function with ‘*fdr*’ method). Genes with a FDR < 1% and |*β_r_*| > 0.2 were considered to be as differentially expressed between populations (i.e., “raw” popDEGs). We adjusted for cellular composition within each lineage *L*, by introducing into model (3) a set of variables (*F_j_*)_*j*∈*l*_ encoding the frequency in the PBMC fraction of each cell type *j* comprising the lineage (e.g., naïve, effector, and regulatory subsets of CD4+ T cells).

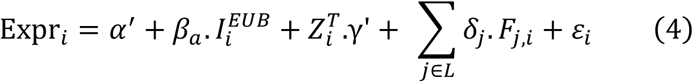

The notation is as above, with *α*’, *β_a_*, *γ*’, the fitted parameters of the model. In this model, *δ_j_* is the effect on gene expression of a 1% increase in cell type *j* and *β_a_* indicates the cell composition-adjusted log_2_fold change difference in expression between AFB and EUB individuals. The significance of *β_a_* was calculated as decribed above, with a *sandwich* estimator of variance and the *coeftest* function. FDR was calculated across all conditions and lineages to yield a set of “cell-composition-adjusted” popDEGs. We assessed the impact of cellular composition on differences in gene expression between populations, by defining Student’s test statistic *T*_Δ*β*_ as follows:

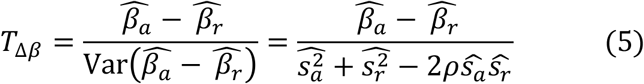

where 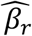 and 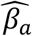 are the raw and cell-composition-adjusted differences in expression between populations, 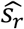 and 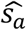 are the estimated standard error of 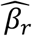 and 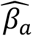 respectively, and *ρ* is the observed correlation in permuted data between the 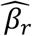 and 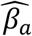 statistics. Under the null hypothesis that population differences are not affected by cellular composition, *T*_Δ*β*_ should follow an approximate Gaussian distribution with mean 0 and variance 1, enabling the definition of a *p*-value *p*_Δ*β*_. We then considered the set of raw popDEGs that (1) were not significant after adjustment (FDR > 1% or |*β_a_*|<0.2) and (2) displayed significant differences between the raw and adjusted effect sizes (|*T*_Δ*β*_ |>1.96) imputable to the effect of cellular composition.

For the assessment of population differences in response to viral stimuli (i.e., popDRGs), we used the same approach, but with the replacement of log-normalized counts with the log-fold change difference in expression between the stimulation conditions for each of the two viruses and non-stimulated conditions.

### Pathway enrichment analyses

We performed functional assessments of the effects of cellular composition variability on differences in gene expression between donors in the basal state and in response to each virus, with the *fgsea* R package (v1.18.1) (ref.^73^) and default options. This made it possible to perform a gene set enrichment analysis with population differences in each lineage ranked by the magnitude of the effect of ancestry on the expression or response of the gene before (β_*r*_) and after (β_*a*_) adjustment for differences in cellular composition.

### Fine mapping of expression quantitative trait loci (eQTL)

For eQTL mapping, we used variants with MAF >5% in at least one of the three populations considered, resulting in a set of 10,711,657 SNPs, of which 4,164,060 were located < 100kb from a gene. We used *MatrixEQTL* (v2.3) (ref.^74^) to map eQTLs in a 100-kb region around each gene and obtain estimates of eQTL effect sizes and their standard error. eQTL mapping was performed separately for each immune lineage/cell type and condition, based on rank transformed gene expression values. eQTL analyses were performed adjusting for population, age, chromosomal sex, cell composition (within each lineage), as well as cell mortality and total number of cells in the sample, and a data-driven number of surrogate variables included to capture unknown confounders and remove unwanted variability. Specifically, for each immune lineage/cell type and condition, surrogate variables were obtained using the *sva* function from the *sva* R package (v3.40.0) (ref.^75^) with option *method= ‘two-steps’*, providing all other covariates as known confounders (*mod* argument). The number of surrogate variables to use in each lineage/cell type and condition was determined automatically based on the results from *num.sv* function with *method= ‘be’* (ref.^75^).

For each gene, immune lineage/cell type and stimulation condition, *Z*-values (i.e., the effect size of each eQTL divided by the standard error of effect size) were then calculated, and the fine mapping of eQTLs was performed with *SuSiE* (v0.11.42) (ref.^76^) (*susie_rss* function of the *susieR* R package), with a default value of up to 10 independent eQTLs per gene. Imputed genotype dosages were extracted in a 100-kb window around each gene and regressed against the population of origin (i.e., AFB, EUB or ASH). Genes with <50 SNPs in the selected window were discarded from the analysis. Pairwise correlations between the population-adjusted dosages were then assessed, to define the genotype correlation matrix to be used for the fine mapping of eQTLs. In rare cases (<0.1% of tested genes × conditions combinations), the *susie_rss* function failed to converge, even when the number of iterations was increased to >10^6^. These runs were, thus, discarded, and the associated eQTLs were assigned a null *Z*-score during FDR computation (see below). For each eQTL, the *index* SNP was defined as the SNP with the highest posterior inclusion probability (i.e., the α parameter in the output of *SuSiE*) for that eQTL, and the 95% credible interval was obtained as the minimal set of SNPs *S* such that *α_s_* > 0.01 for all *s* ∈ *S* and ∑_*s*∈*s*_ α_*s*_ > 0.95. Only eQTLs with a log-Bayes factor (*lbf*) > 3 were considered for further analyses.

For each lineage and set of stimulation conditions, each eQTL identified by *SuSiE* was assigned an *eQTL evidence score*, defined as the absolute *Z*-value of association between the eQTL index SNP and the associated gene. We then used a pooled permutation strategy to define the genome-wide number of significant eQTLs (i.e., eQTL × gene combinations) expected under the null hypothesis, for different thresholds of the *eQTL evidence score*. We repeated the eQTL mapping procedure on the dataset after randomly permuting genotype labels within each population. We then counted, for each possible *evidence score* threshold *T*, the number of eQTLs identified in the observed and permuted data. Finally, we retained as a significance threshold the lowest threshold giving a number of significant eQTLs in the permuted data (false positives) of less than 1% the number of eQTLs identified in the observed data (false positives + true positives).

### Aggregation of eQTLs across cell types and stimulation conditions

The eQTL index SNP may differ between cellular states (immune lineage/cell type and stimulation condition), even in the presence of a single causal variant. It is therefore necessary to aggregate eQTLs to ensure that the same locus is tagged by a single variant across cellular states. To this end, we applied the following procedure, for each gene:

1. Let *C_g_* be the set of cellular states where a significant eQTL was detected for gene *g*, and *S_g_* be the associated list of eQTLs (i.e., cellular state × index SNPs). We aim to define a minimal set of SNPs, *M_g_*, that overlaps the 95% credible intervals of all significant eQTLs in *S_g_*.
2. For each SNP *s* in a 100-kb window around each gene, compute the expected number of cellular states where the SNP has a causal effect on gene expression *E*[*N_c_*(*s*)] as:

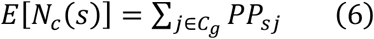

where *PP_sj_* is the posterior probability that SNP *s* has a causal effect on the expression of gene *g* in the cellular state *j* (cell type × condition).
3. Find the SNP *s* that maximizes *E* [*N_c_*(*s*), and add it to *M_g_*
4. Remove from *S_g_*, all eQTLs where the 95% credible interval contains SNP *s*
5. Repeat steps 1-3 until *S_g_* is empty.

At the end of this procedure, *M_g_* provides the list of independent eQTL index SNPs (referred to as eSNPs) for gene *g*, for which we extracted summary statistics across all cellular states.

### Mapping of response eQTLs

For the mapping of response eQTLs (reQTLs), we repeated the same procedure as for the mapping of eQTLs, using rank-transformed log_2_ fold change as input rather than gene expression. This included reQTL mapping with *MatrixEQTLL*^74^, fine mapping with *SuSiE*^76^, permutation-based FDR computation, and aggregation of reQTL across immune lineages, cell types and stimulation conditions. Surrogate variables were computed directly from log_2_fold changes. For IAV-infected monocytes (detected only in the IAV condition), fold changes were computed relative to CD14^+^ monocytes in the non-stimulated condition. This produced a list of independent reQTL index SNPs *M*’, similar to that obtained for eQTLs, for which we extract summary statistics across all cellular states.

### Sharing of eQTLs across cell types and stimulation conditions

After extracting the set *M* of independent eSNPs across all genes, we defined ‘cell-type-specific eQTLs’ as eQTLs significant genome-wide in a single cell type. We accounted for the possibility that some eQTLs may be missed in specific cell types due to a lack of power, by introducing a second definition of eQTL sharing based on nominal *p*-values. Specifically, we considered an eQTL to be cell type-specific at a nominal significance if, and only if, it was significant genome-wide in a single cell type and its nominal *p*-value of association was greater than 0.01 in all other cell types. For each pair of cell types, the correlation of eQTL effect sizes was calculated on the set of all eQTLs passing the nominal significance criterion (*p* < 0.01) in at least one of the two cell types. To understand how the effect of genetics on immune response varies between SARS-CoV-2 and IAV, we defined an interaction statistic that enabled us to test for differences in reQTL effect size between the two viruses. Specifically, within each immune lineage/cell type, we defined:

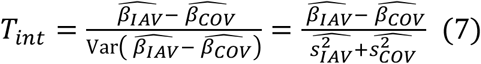

When the reQTL effect size is identical between the two viruses, we expect *T_int_* to be normally distributed around 0 with variance 1, allowing to derive an interaction *p*-value. We thus defined as *virus-dependent reQTLs* those with a nominal interaction *p*-value < 0.01 and as *virus-specific reQTLs* those that passed a nominal *p*-value threshold of 0.01 in only one of the two stimulation conditions.

### Mediation analyses

For all popDEGs and popDRGs, we evaluated the proportion of the difference in expression or response to viral stimulation between populations attributable to either genetic factors (i.e., eQTLs) or cellular composition, with the *mediate* function of the *mediation* package of R (v4.5.0) (ref.^77^). Mediation analysis made it possible to separate the differences in expression/response between populations that were mediated by genetics (i.e., differences in allele frequency of a given eQTL between populations, *ζ_g_*), or cellular composition (i.e., difference in cell type proportions between populations, *ζ_c_*) from those occurring independently of the eQTL/cell type considered (independent or direct effect *δ*). It was then possible to estimate the respective proportion of population differences mediated by genetics *τ_g_* and cellular composition *τ_c_* as 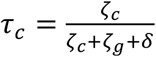 and 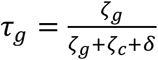 with *ζ_c_* + *ζ_g_* + δ corresponding to the total differences in expression/response between populations. For each popDEG and popDRG, we focused on either (i) the most strongly associated SNP in a 100-kb window around the gene, regardless of the presence or absence of a significant (r)eQTL, or (ii) the cell type differing most strongly between populations in each lineage (i.e., CD16^+^ monocytes in the myeloid lineage, *κ* light chain-expressing memory B cells in the B-cell lineage, effector cells in CD4^+^ T cell lineage, CD8+ EMRA cells in the CD8^+^ T-cell lineage, and memory cells in the NK cell lineage). For each popDEG and potential mediator M, we then ran *mediate* with the following models:

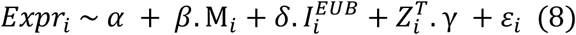

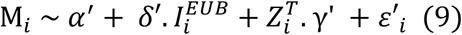

where *Expr_i_* corresponds to normalized expression values in the cell type/condition under consideration, *α* and *α*’ are two intercepts, *β* is the effect of the mediator M_*i*_ on gene expression, *δ* and *δ*’ are the (direct) effect of population (captured through the indicator variable 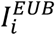) on gene expression and on the mediator, *γ* and *γ*’ capture the confounding effect of covariates (i.e. age, and cell mortality) on both gene expression and the mediator, and *ε*_*i*_ and *ε*’_*i*_ are normally distributed residuals. For popDRGs, we used the same approach, replacing normalized gene expression values with the log_2_fold change in gene expression between the stimulated and unstimulated states.

### Detection of signals of natural selection

We avoided SNP ascertainment bias, by performing natural selection analyses with high-coverage sequencing data from the 1,000 Genomes (1KG) Project^78^. We downloaded the GRCh38 phased genotype files from the New York Genome Center FTP server and calculated the pairwise *F*_ST_ (ref.^79^) between our three study populations (AFB, EUB, or ASH) and all 1KG populations, to identify the 1KG populations who were the most genetically similar to our study populations. All selection and introgression analyses (see section ‘Archaic introgression analyses’) were based on the Yoruba from Ibadan, Nigeria (YRI), Utah residents with Northern and Western European ancestry (CEU) and Southern Han Chinese (CHS) populations, as these 1KG populations had the lowest *F*_ST_ values with our three study groups. We filtered the data to include only autosomal bi-allelic SNPs and insertions/deletions (indels), and removed sites that were invariant (i.e., monomorphic) across the three populations. We identified loci presenting signals of positive selection (local adaptation) with the population branch statistic (PBS)^38^, based on the Reynold’s *F*_ST_ estimator^79^ between pairs of populations. PBS values were calculated for the YRI, CEU, and CHS populations separately, with the other two populations used as the control and outgroup. For each population, genome-wide PBS values were then ranked, and variants with PBS values within the top 1% were considered putative targets of selection. For annotation of the selected eQTLs, the ancestral and derived states at each site were inferred from six-way EPO multiple alignments for primate species (EPO6, available from ftp://ftp.ensembl.org/pub/release-71/emf/ensembl-compara/epo_6_primate/), and the effect size was reported for the derived allele. For sites without an ancestral/derived state in the EPO6 alignment, the effect of the allele with the lowest frequency worldwide was reported.

We assessed the extent to which different sets of eQTLs displayed signals of local adaptation in permutation-based enrichment analyses. For each population, we compared the mean PBS values at (r)eQTLs for each set of cell type/stimulation condition with the mean PBS values obtained for 10,000 sets of randomly resampled sites. Resampled sites were matched with eQTLs for minor allele frequency (mean MAF across the three populations, bins of 0.01), LD scores (quintiles), and distance to the nearest gene (bins of 0-1 kb, 1 kb-5 kb, 10 kb-20 kb, 20 kb-50 kb, >100 kb). For each population and set of eQTLs, we defined the fold-enrichment (FE) in positive selection as the ratio of observed/expected values for mean PBS and extracted the mean and 95% confidence interval of this ratio across all resamplings. One-sided resampling *p*-values were calculated as the number of resamplings with a FE>1 divided by the total number of resamplings. Resampling *p*-values were then adjusted for multiple testing by the Benjamini-Hochberg method.

### Detecting and dating episodes of local adaptation

We inferred the frequency trajectories of all eQTLs and reQTLs during the past 2,000 generations (i.e., 56,000 years before the present, with a generation time of 28 years), systematically by using *CLUES* (commit no. 7371b86, 27 may 2021) (ref.^39^). We first used *Relate* (v1.1.8) (ref.^80^) on each population separately, to reconstruct tree-like ancestral recombination graphs (ARGs) around each SNP in the genome and to estimate effective population sizes across time based on the rate of coalescence events over the inferred ARGs. Using *CLUES*^39^, we then estimated at each eQTL or reQTL, the most likely allele frequency trajectories for each sampled ARG and averaged these trajectories across all possible ARGs.

We then analyzed changes in inferred allele frequencies over time, to identify selection events characterized by a rapid change in allele frequency. We considered the posterior mean of allele frequency at each generation and smoothed the inferred allele frequency trajectories by *loess* regression to ensure progressive changes in allele frequency over time, and to minimize the artifacts induced by the inference process. Finally, for each variant and in each population, we calculated the change in allele frequency *f* at each generation as the difference in the smoothed allele frequency between two consecutive generations:

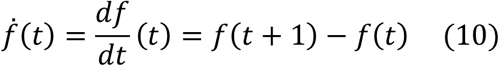

Under an assumption of neutrality, the count of a particular allele at generation *t*+1 is the result of a Bernoulli trial parameterized *B*(*Nf*), where *N* is the size of the haploid population. The variance of allele frequencies at generation *t*+1 is, therefore, greater for alleles present at higher frequencies in generation *t*,

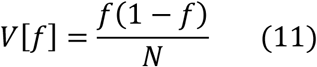

We adjusted for this, by scaling the change in allele frequency 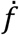 by a normalizing factor dependent on the allele frequency at generation *t*, such that the variance of the normalized change in allele frequency 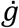 was the same across all variants,

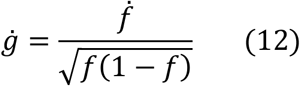

Finally, at each generation, we divided the normalized change in allele frequency 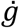 by its standard deviation across all eQTLs and reQTLs, to calculate a *Z*-score for detecting alleles for which the normalized change in allele frequency exceeded genome-wide expectations,

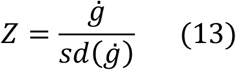

For each variant and generation, we then considered an absolute *Z*-score > 3 to constitute evidence of selection and we inferred the onset of selection of a variant as the first generation in which |*Z*| > 3.

### Simulations, power, and type I error estimates

We assessed the ability of our approach to detect (and date) events of natural selection correctly from the trajectories of allele frequencies, by using simulations with SLiM (v4.0.1) (ref.^81^) under various selection scenarios. Simulations were performed under a Wright-Fisher model for a single mutation occurring ~5000 generations ago, at a frequency varying from 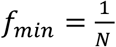 to 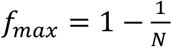 in steps of 1%, where *N* is the simulated population size. We allowed population size to vary over time according to published estimates^80^ for the YRI, CEU and CHS populations (Supplementary Fig. S9b). We then performed simulations both under an assumption of neutrality (1000 simulations for each starting frequency) and assuming a 200 generations-long episode of selection (100 simulations for each starting frequency and selection scenario). Selection episodes were simulated with an onset of selection 1000, 2000, 3000, or 4000 generations ago, and with a selection coefficient ranging from 0.01 to 0.05 (Supplementary Fig. S9c). We saved computation time, by performing a 10-fold scaling in line with SLiM recommendations. For each selected scenario and variant, simulated allele frequencies were retrieved every 10 generations, and smoothed with the *loess* function of R using default parameters. We then calculated normalized differences in smoothed allele frequencies for each simulated variant and scaled these differences at each generation, based on their standard deviation among neutral variants, to obtain *Z*-scores. For each selection scenario, we focused on the center of the selection interval and determined the type I error and power for various thresholds of absolute *Z*-scores varying from 0 to 6. We found that a threshold of 3 yielded both a low type I error (<0.2% false positives) and a satisfactory power for detecting selection events (Supplementary Fig. S9c). Finally, at each generation, we estimated the percentage of simulations, under an assumption of neutrality or a particular selection scenario, for which the absolute *Z*-score exceeded a threshold of 3. We found that significant *Z*-scores were equally rare at each generation under the assumption of neutrality, but that selected variants presented a clear and localized enrichment in significant *Z*-scores for intervals in which we simulated selection (Supplementary Fig. S9d).

### Archaic introgression analyses

For the definition of regions of the modern human genome of archaic ancestry (Neanderthal or Denisovan), we downloaded the VCFs from the high-coverage Neanderthal Vindija^82^ and Denisovan Altai^83^ genomes (human genome assembly GRCh37; http://cdna.eva.mpg.de/neandertal/Vindija/) and applied the corresponding genome masks (FilterBed files). We then removed sites within segmental duplications and lifted over the genomic coordinates to the GRCh38 assembly with *CrossMap* (v0.6.3) (ref.^84^). We used two statistics to identify introgressed regions in the CEU and CHS populations: (i) conditional random fields (CRF)^85,86^, which uses reference archaic and outgroup genomes to identify introgressed haplotypes; and (ii) the S’ method^87^, which identifies stretches of probably introgressed alleles without requiring the definition of an archaic reference genome.

For CRF-based calling, we phased the data with *SHAPEIT4* (v4.2.1) (ref.^55^), using the recommended parameters for sequence data, and focused on bi-allelic SNPs for which the ancestral/derived state was unambiguously defined. We then performed two independent runs of CRF to detect haplotypes inherited from Neanderthal or Denisova. For Neanderthal-introgressed haplotypes, we used the Vindija Neanderthal genome as the archaic reference and YRI individuals merged with the Altai Denisovan genome as the outgroup. For Denisovan-introgressed haplotypes, we used the Altai Denisovan genome as the archaic reference panel and YRI individuals merged with the Vindija Neanderthal genome as the outgroup. Results from the two independent CRF runs were analyzed jointly, and we retained alleles with a marginal posterior probability *P*_Neanderthal_ ≥ 0.9 and *P*_Denisova_ < 0.5 as Neanderthal-introgressed haplotypes and those containing alleles with *P*_Denisova_ ≥ 0.9 and *P*_Neanderthal_ < 0.5 as Denisovan-introgressed haplotypes. For the S’-based calling of introgressed regions, we considered all biallelic SNPs with an allele frequency < 1% in the YRI population to be Eurasian-specific alleles. We then ran the *Sprime* software (v.07Dec18.5e2) (https://github.com/browning-lab/sprime) separately for the CEU and CHS populations, to identify and score putatively introgressed regions containing a high density of Eurasian-specific alleles. Putatively introgressed regions with a S’ score >150,000 were considered to be introgressed. This cutoff score has been shown to provide a good trade-off between power and accuracy based on simulations of introgression under realistic demographic scenarios^87^. For both calling methods (i.e., CRF and S’), we used the recombination map from the 1,000 Genomes (1KG) Project Phase 3 data release^51^.

After the calling of introgressed regions throughout the genome for each population, we defined SNPs of putative archaic origin (archaic SNPs, aSNPs) as those (i) located in an introgressed region defined by either the CRF or S’ method, (ii) with one of their alleles being rare or absent (MAF < 1%) in the YRI population, but present in the Vindija Neanderthal or Denisovan Altai genomes, and (iii) in high LD (*r*^2^ > 0.8) with at least two other aSNPs and, to exclude incomplete lineage sorting, comprising an LD block of > 10 kb. This yielded a set of 100,345 high-confidence aSNPs (Supplementary Table S7a). We further categorized aSNPs as of *Neanderthal origin, Denisovan origin* or *shared origin* according to their presence/absence in the Vindija Neanderthal and Denisovan Altai genomes. Finally, we considered any site that was in high LD with at least one aSNP in the same population in which introgression was detected to be *introgressed*, and classified introgressed haplotypes as of *Neanderthal origin, Denisovan origin*, or *shared* origin according to the most frequent origin of aSNPs in the haplotype. For introgressed SNPs, we defined the introgressed allele as (i) the allele rare or absent from individuals of African ancestry if the SNP was an aSNP, and (ii) for non-aSNPs, the allele most frequently segregating with the introgressed allele of linked aSNPs. In each population, introgressed alleles with a frequency in the top 1% for introgressed alleles genome-wide were considered to present evidence of *adaptive introgression*.

The enrichment of introgressed haplotypes in eQTLs or reQTLs was assessed separately for each population (CEU and CHS), first by stimulation condition and then by cell type within each condition. Within each cell type/stimulation condition, we considered the set of all (r)eQTLs for which the index SNP displayed at least a marginal association (Student’s *p* < 0.01) with gene expression. For each population and (r)eQTL set, we then grouped (r)eQTLs in high LD (*r*^2^ > 0.8), retaining a single representative per group, and counted the total number of (r)eQTLs for which the index SNP was in LD (*r*^2^ > 0.8) with an aSNP (i.e., introgressed eQTLs). We then used *PLINK (v1.9) --indep-pairwise* (with a 500-kb window, 1 kb step, an *r*^2^ threshold of 0.8, and a MAF > 5%) (ref.^52^), to define tag-SNPs for each population, and we determined the expected number of introgressed SNPs by resampling tag-SNPs at random with the same distribution for MAF, LD scores, and distance to the nearest gene. We performed 10,000 resamplings for each (r)eQTL set and population. One-sided resampling-based *p*-values were calculated as the frequency at which the number of introgressed SNPs among resampled SNPs exceeded the number of introgressed SNPs among (r)eQTLs. Resampling-based *p*-values were then adjusted for multiple testing by the Benjamini–Hochberg method.

We searched for signals of adaptive introgression, by determining whether introgressed haplotypes that altered gene expression were introgressed at a higher frequency than introgressed haplotypes with no effect on gene expression. For each stimulation cell type/condition, we focused on the set of introgressed eQTLs segregating with a MAF > 5% in each population (retaining a single representative per LD group) and compared the frequency of the introgressed allele with that of introgressed tag-SNPs genome-wide. We modeled *r*_(*Freq*)_, the (rank-transformed) frequency of introgressed tag-SNPs according to the presence/absence of a linked eQTL 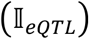, and the mean MAF of the SNP across the three populations (giving a higher power for eQTL detection).

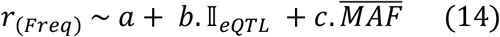

where 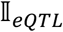 is an indicator variable equal to 1 if the SNP is in LD with an eQTL (*r*^2^>0.8) and 0 otherwise, 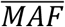 is the mean MAF calculated separately for each population, *a* is the intercept of the model, *b* measures the difference in rank *r_Freq_* between eQTLs and non eQTLs, and *c* is a nuisance parameter capturing the relationship between 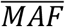 and *r*_(*Freq*)_ Under this model, the difference in frequency between eQTLs and non-eQTLs can be tested directly in a Student’s *t* test with *H*_0_: *b* = 0.

### Enrichment in COVID-19-associated loci and colocalization analyses

We downloaded summary statistics from the COVID-19 Host Genetics Initiative (release 7: https://www.covid19hg.org/results/r7) (ref.^7^) for three GWAS: (i) A2 - very severe respiratory cases of confirmed COVID-19 *vs*. the general population; (ii) B2 - hospitalized COVID-19 cases *vs*. the general population; (iii) C2 – confirmed COVID-19 cases *vs*. the general population. We assessed the enrichment in eQTLs and reQTLs of COVID-19 susceptibility/severity loci by considering, for each eQTL/reQTL, the A2, B2 or C2 *p*-values of the index SNP and calculating the percentage of eQTLs/reQTLs with a significant GWAS *p*-value of 10^-4^. This percentage was then compared to that obtained for the resampled set of SNPs, matched for distance to the nearest gene (bins of 0-1, 1-5, 5-10, 10-20, 20-50, and 50-100 kb) and MAF (1% MAF bins). We performed 10,000 resamplings for each set of eQTLs/reQTLs tested. The use of different *p*-value thresholds for COVID-19-associated hits (10^-3^ to 10^-5^) yielded similar results

To identify specific eQTLs/reQTLs colocalized with GWAS hits, we first considered all (r)eQTLs for which the index SNPs were located within 100 kb of a SNP associated with COVID-19 susceptibility/severity (*p*-value < 10^-5^). For each immune lineage/cell type, and condition for which the eQTL/reQTL reached genome-wide significance, we next extracted all SNPs in a 500-kb window around the index SNP for which summary statistics were available for both the eQTLs/reQTLs and COVID-19 GWAS phenotypes (A2, B2, and C2) and performed a colocalization test using the *coloc.signals* function of the *coloc* (v5.1.0) package of R. We set a prior probability for colocalization *p_12_* of 10^-5^ (i.e., the recommended default value). Any pair of eQTL or reQTL/COVID-19 phenotypes with a posterior probability for colocalization PP.H4 > 0.8 was considered to display significant colocalization.

### Statistical Analyses

Unless explicitly specified, all statistical tests are two-sided and based on measurements from independent samples.

## Data availability

The RNA sequencing data generated and analyzed in this study have been deposited in the Institut Pasteur data repository, OWEY, which can be accessed via the following link: https://doi.org/XXXX. The genome-wide genotyping data generated or used in this study have been deposited in OWEY and can be accessed at the following URL: https://doi.org/XXXX. Data access and use is restricted to academic research related to the variability of the human immune response.

## Code availability

All custom computer code or algorithms used in this study are available from github (https://github.com/h-e-g/popCell_SARS-CoV-2).

## Inclusion and Ethics

The current research project builds on samples collected in Ghent (Belgium) and Hong-Kong SAR (China) and has been conducted in collaboration with local researchers. Roles and responsibilities were agreed amongst collaborators ahead of the research. Research conducted in this study is relevant to local participants and has been reviewed by local ethics committees (committee of Ghent University, Belgium, n° B670201214647; Institutional Review Board of the University of Hong-Kong; n° UW 20-132), and the relevant French authorities (CPP, CCITRS and CNIL). This study was also monitored by the Ethics Board of Institut Pasteur (EVOIMMUNOPOP-281297). All manipulations of live viruses were performed in a high security BSL-3 environment.

## Acknowledgments

We thank all members of the Human Evolutionary Genetics Laboratory, and Bjorn-Axel Olin in particular, and Monique Van der Wijst, Dylan de Vries and Lude Franke for helpful discussions, Leo Speidel for help in running *Relate* and interpreting the results, and Anja Coen and Frederic Clément for assistance with sample collection. The Human Evolutionary Genetics Laboratory is supported by the *Institut Pasteur*, the *Collège de France*, the *Centre Nationale de la Recherche Scientifique* (CNRS), the *Agence Nationale de la Recherche* (ANR) grants COVID-19-POPCELL (ANR-21-CO14-0003-01), POPCELL-REG (ANR-22-CE12-0030-01) and COVIFERON (ANR-21-RHUS-0008), the EU HORIZON-HLTH-2021-DISEASE-04-07 grant UNDINE (no. 101057100), the French Government’s *Investissement d’Avenir* program, *Laboratoires d’Excellence* “Integrative Biology of Emerging Infectious Diseases” (ANR-10-LABX-62-IBEID) and *“Milieu Intérieur”* (ANR-10-LABX-69-01), the *Fondation pour la Recherche Médicale* (Equipe FRM DEQ20180339214), the *Fondation Allianz-Institut de France*, and the *Fondation de France* (no. 00106080). K.L., J.T.K.W. and M.P. are supported by the Health and Medical Research Fund Commissioned Research on the Novel Coronavirus Disease, Hong Kong SAR (COVID190126), S.A.V by the Theme-based Research Scheme of the Research Grants Council of the Hong Kong SAR (T11-705/21-N, SAV: T11-712/19-N), and M.P., R.B. and D.D. by *InnoHK*, an initiative of the Innovation and Technology Commission, the Government of the Hong Kong SAR.

## Author Contributions

M.R. and L.Q.-M. conceived and supervised the study; Y.A., A.B., M.O’N. and M.R. designed experiments; Y.A., A.B., Z.L., M.O’N. and S.H.M. conducted the experiments; Y.A., M.O’N, J.M.-R., G.K. and M.R. designed and performed computational analyses; V.B. conducted the SIMOA experiments; G.L.-R., C.-K.L., K.L., J.T.K.W., M.P., R.B. and S.A.V. managed clinical protocols and recruited donors; N.S., G.B.-S., S.P. and O.S. obtained SARS-CoV-2 strains and helped to design the stimulation experiments; C.C., M. M., M.H., V.L., F.L, R.P.-R., L.A., J-L.C., D.D and E.P. advised on experiments and data analyses and interpretation; Y.A., A.B., M.R. and L.Q.-M. interpreted the data and wrote the manuscript, with critical input from all authors.

---

**Supplementary Figure 1.**
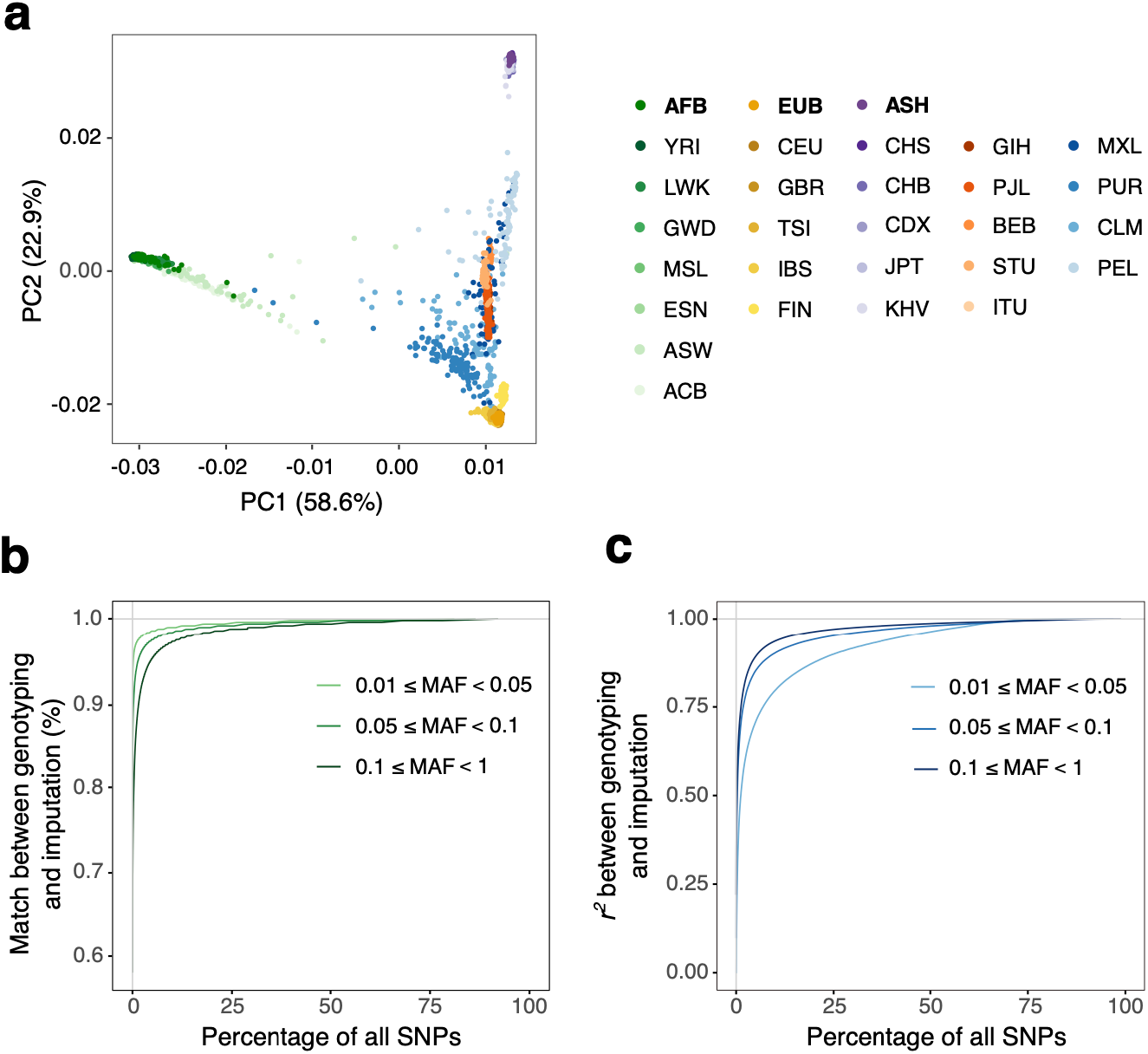
Genetic structure of study populations and SNP imputation. **a,** Principal component analysis of genotyping data. Each dot corresponds to an individual. Study samples (AFB, EUB, and ASH, in bold typeface) are projected jointly with 1,000 Genomes populations of various ancestries including African (dots colored in green gradients), European (dots colored in yellow gradient), East Asian (dots colored in purple gradient), South Asian (dots colored in orange gradient) and American (dots colored in blue gradients). Abbreviations for each individual population can be found in ref.^51^. **b** and **c,** Quality control of genotype imputation. Distribution of *r*^2^ between genotyped and imputed SNPs (**b**) and genotype accuracy (**c**) obtained by 100-fold cross-validation and shown separately for different bins of minor allele frequency (MAF).

**Supplementary Figure 2.**
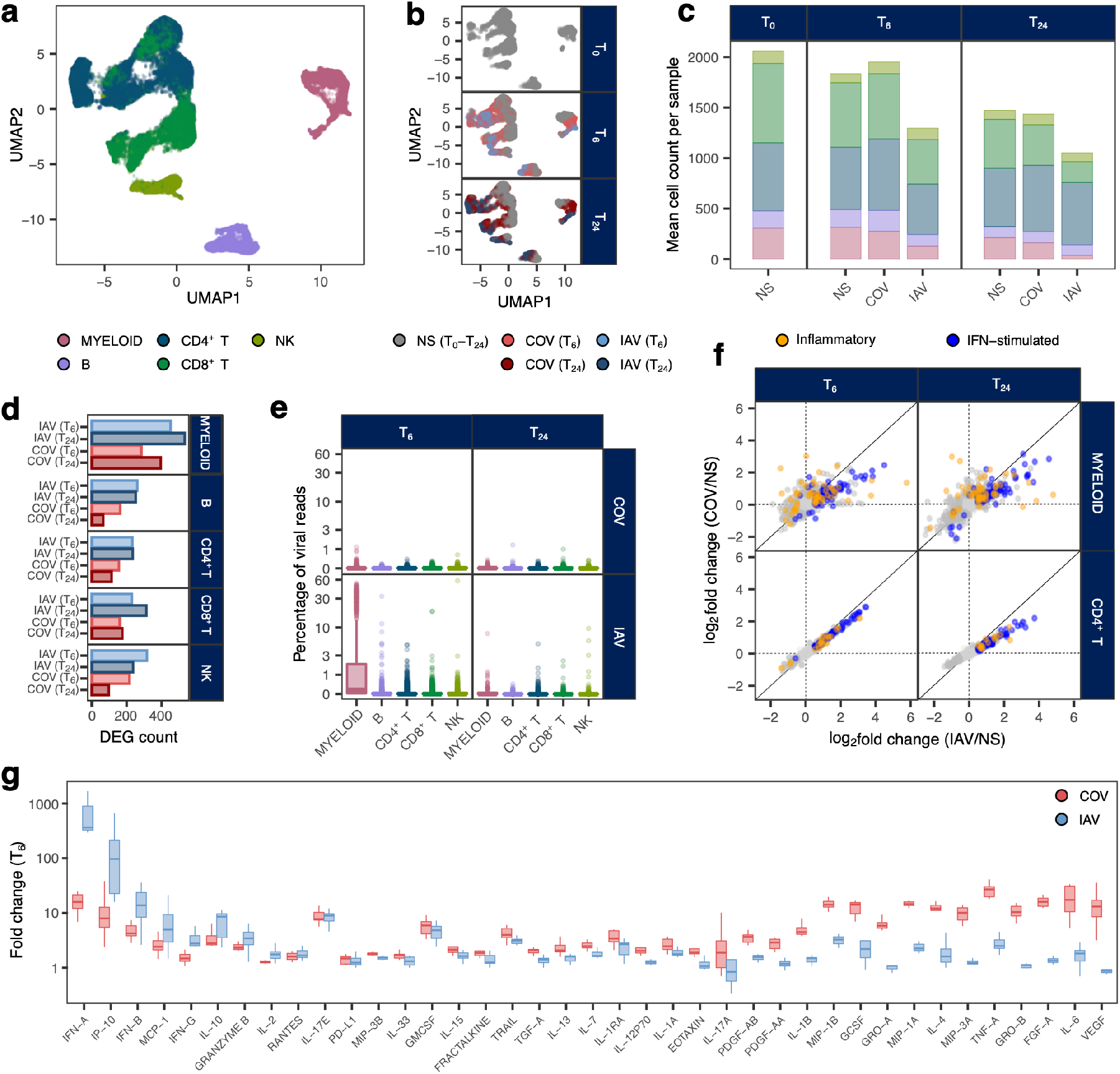
Single-cell kinetics of the immune response to RNA viruses. **a, b,** Uniform manifold approximation and projection (UMAP) of 86,363 peripheral blood mononuclear cells (PBMCs), mock-stimulated (NS) or stimulated with SARS-CoV-2 (COV) or influenza A virus (IAV) for 0, 6 or 24 hours. **c,** Mean cell-type counts per individual, set of stimulation conditions and time point. **d,** Number of differentially expressed genes (DEG; absolute log_2_ fold change (|log_2_FC|) > 0.5, FDR < 0.01) after 6 or 24 hours of stimulation with SARS-CoV-2 or IAV relative to non-stimulated controls. **e,** Percentage of reads mapping to the SARS-CoV-2 or IAV genomes per cell, after 6 or 24 hours of stimulation, split by major immune lineage. **f,** Comparison of inflammatory and interferon-stimulated transcriptional responses of myeloid cells and CD4^+^ T cells (as an example of a lymphoid cell type) after 6 or 24 hours of stimulation with SARS-CoV-2 or IAV. **g,** Cytokine protein responses to 6 hours of stimulation with SARS-CoV-2 or IAV. In **a, c** and **e,** the colors indicate the immune lineages inferred from single-cell transcriptome data. In **b**, **d** and **g,** the colors indicate the stimulation condition and time post-stimulation. In **e** and **g**, boxplots are defined as follows: middle line, median; box limits, upper and lower quartiles; whiskers, 1.5× interquartile range; points, outliers.

**Supplementary Figure 3.**
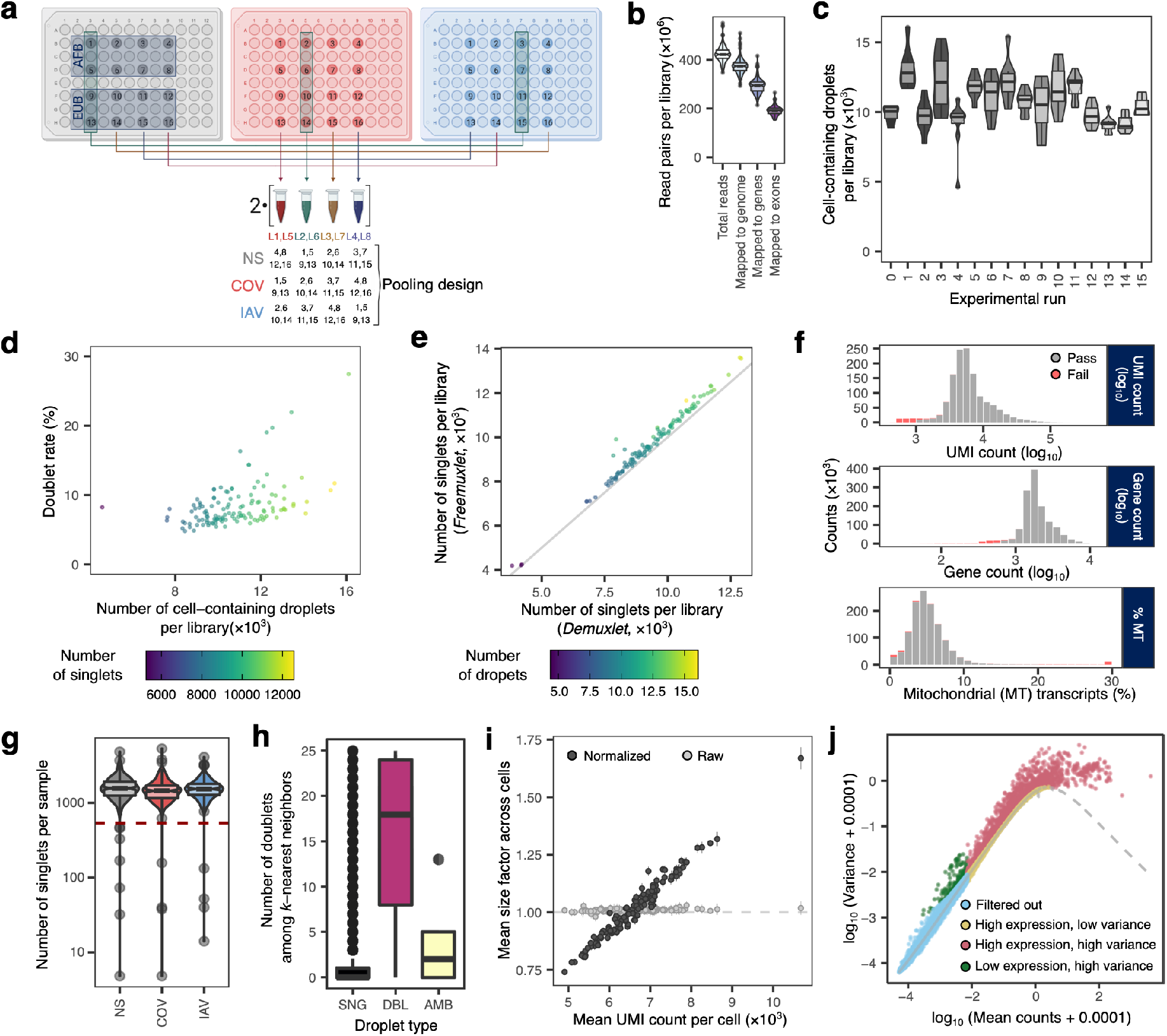
Quality control of single-cell RNA-seq data. **a**, Experimental design. During each experimental run, PBMCs from 16 individuals (numbered 1 to 16) were processed in three different sets of experimental conditions (colored plates), and the resulting samples were split into four pools of 12 samples (four non-stimulated (NS), four influenza A virus-stimulated (IAV), four SARS-CoV-2-stimulated (COV)). Each pool was then processed on two independent libraries to increase the number of cells per sample (eight pools of 12 samples in total). **b,** Library sequencing depth, and total number of reads aligned either genome-wide, in the genic region, or over the coding exons. **c,** Distribution of the number of cell-containing droplets detected per library across the 16 experimental runs performed. **d,** Percentage of doublets per library as a function of the number of cell-containing droplets detected. Colors reflect the inferred number of singlets in the library. **e,** Number of singlets per library inferred with two independent demultiplexing algorithms: *Demuxlet* (supervised) and *Freemuxlet* (unsupervised). Colors reflect the total number of droplets in the library. **f**, Distribution of cells along quality-control metrics in our data set (i.e., UMI count, gene count, and percentage of mitochondrial reads). Cells that were excluded by our hard-threshold filtering are highlighted in red. **g**, Number of high-quality cells per sample (individual × condition), split by stimulation condition. Individuals with <500 cells in at least one sample were excluded (dashed red line). **h**, Number of genetic doublets among k nearest neighbors as a function of the droplet status assigned by *Demuxlet* (SNG: singlet, DBL: doublet, AMB: ambiguous). **i**, Per-library mean of raw and batch-normalized size factors, as a function of the mean number of UMIs per cell in the library. Vertical bars indicate 95% CI of the mean. After normalization with *multiBatchNorm*, size factors successfully capture differences in read depth across libraries. **j**, Filtering of weakly expressed and low-variability genes. For each gene, the variance and mean counts are shown on a log scale (with an offset of 10^-4^). The dashed line indicates the expected relationship between mean and variance under a Poisson distribution (technical noise). Genes are colored according to their expression levels (highly expressed: mean > 0.01) and estimated biological variance (highly variable: biological variance > 0.001). Genes that are both weakly expressed and of low variability (light blue) are excluded from downstream analyses. In **b, c, g,** and **h**, boxplots are defined as follows: middle line, median; box limits, upper and lower quartiles; whiskers, 1.5× interquartile range; points, outliers.

**Supplementary Figure 4.**
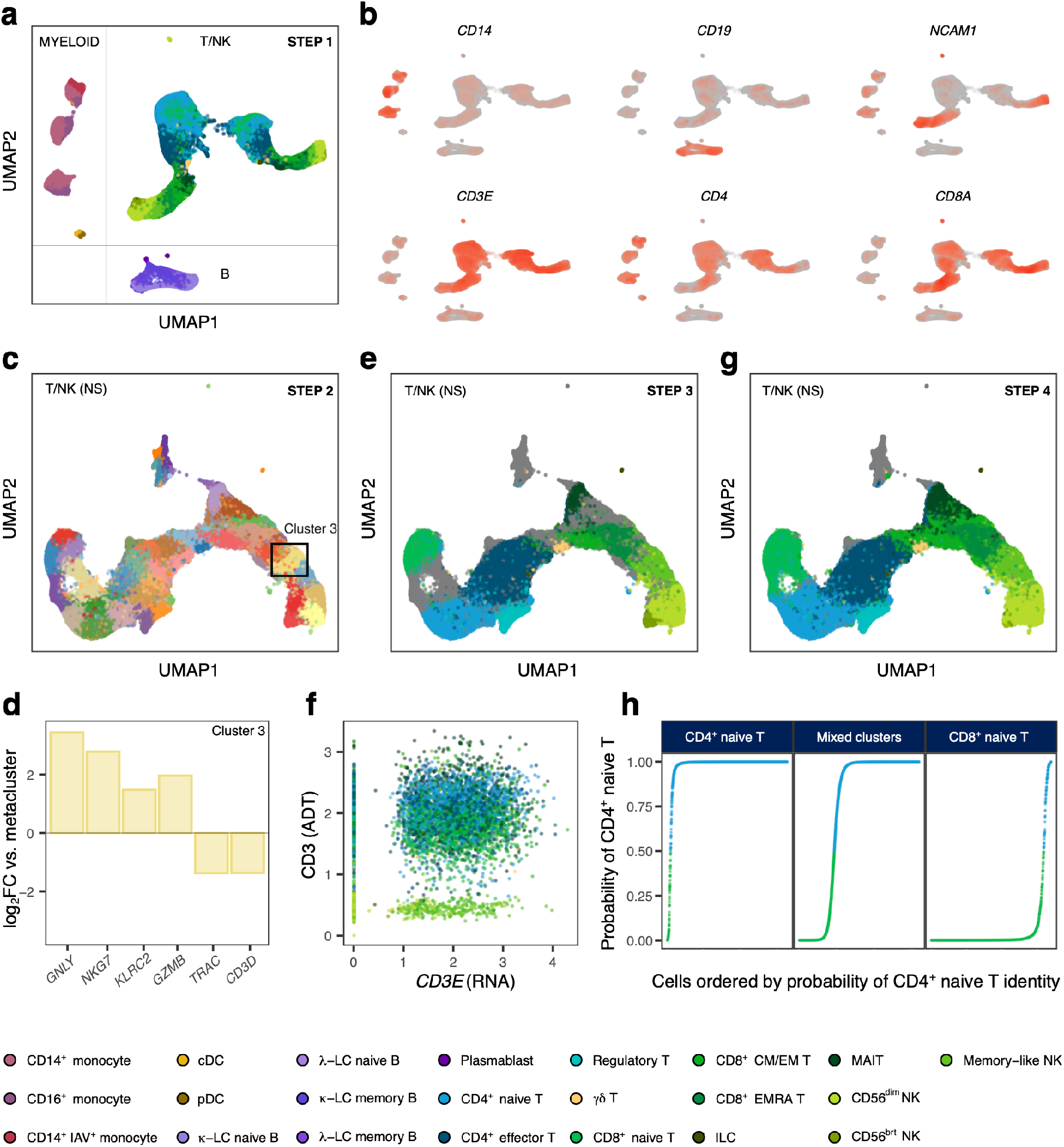
Cell-type assignment according to cluster-based transcriptional profiles and surface protein markers. **a,** Uniform manifold approximation and projection (UMAP) of 1,047,824 PBMCs, either resting (mock-stimulated) or stimulated with SARS-CoV-2 (COV) or influenza A virus (IAV) for 6 hours. **b,** Normalized single-cell RNA UMI count distributions of 6 canonical marker genes. **c,** Graph-based subclustering of the non-stimulated T/NK meta-cluster; cluster 3, initially defined as part of a larger cluster of mixed NK and CD8^+^ T cells, is highlighted. **d,** Log_2_-fold change difference in expression of markers defining NK cell identity between cluster 3 and the rest of the T/NK metacluster. **e,** Cell-type inference based on canonical marker expression in sub-clusters increases the resolution of cell-type identities, but some mixed-identity and unidentified clusters remain (gray). **f,** At the transcriptional level, most of the cells in cluster 3 are *CD3E*-positive, and, thus, associated with lymphocyte lineages, but CITE-seq data show that most cluster 3 cells do not express CD3 protein, hence their assignment to the NK lineage. **g,** Cell-type inference after CITE-seq-based assignment and resolution of mixed-identity clusters by linear discriminant analysis (LDA). Unassigned cells (gray) are discarded. **h,** Assignment of cells from mixed-identity clusters, based on previously identified clusters. In this example, LDA models are trained on data from 10,000 confidently identified naive CD4^+^ and naive CD8^+^ T cells, making it possible to assign most cells from a mixed cluster to one of the two target identities.

**Supplementary Figure 5.**
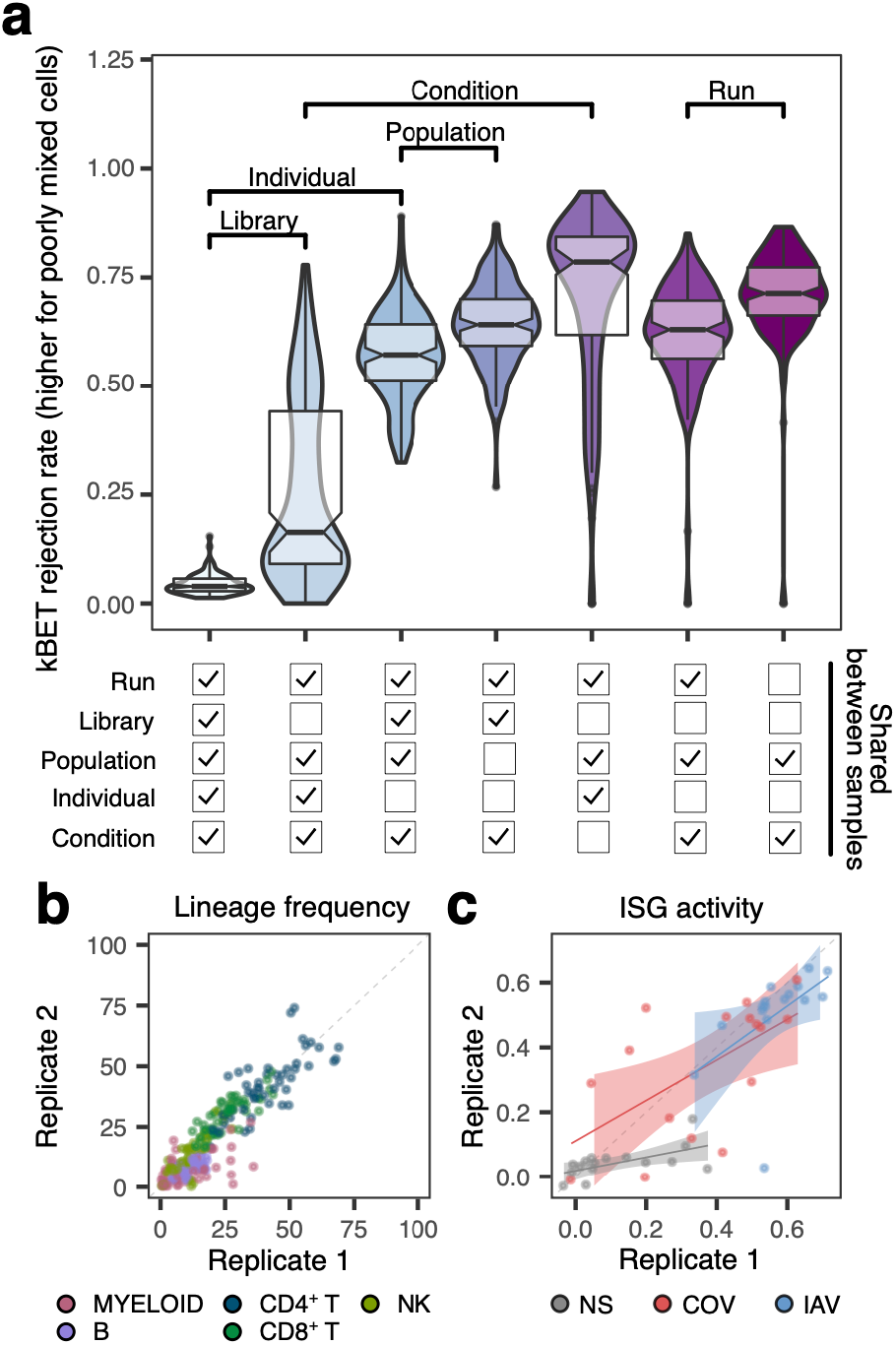
Batch effects and replicability of single-cell experiments. **a,** Effect of technical and biological variation on the clustering of cells. For each comparison, we consider a random subset of 150 pairs of samples (run × library × individual × condition). For each pair of samples, violins and boxplots show the distribution of the kBET rejection rate, which increases when cells from different samples tend to cluster separately (middle line: median; box limits: upper and lower quartiles; whiskers: 1.5× interquartile range; points: outliers). For self-comparisons, cells from the same sample were randomly split into two groups before kBET calculation. Comparisons for quantifying the effects of various factors (e.g., run, library preparation, individual, population or stimulation condition) on cell mixing are highlighted. For all comparisons shown, differences in kBET are significant for a Wilcoxon rank-sum test *p-*value < 10^-9^. **b** and **c,** Comparison of estimated lineage proportions (b) and mean ISG expression (c) between replicate samples processed separately across different runs (cells from the same individual thawed on a different day and stimulated in the same conditions). In **c**, regression lines and the 95% confidence interval are shown for each stimulation condition.

**Supplementary Figure 6.**
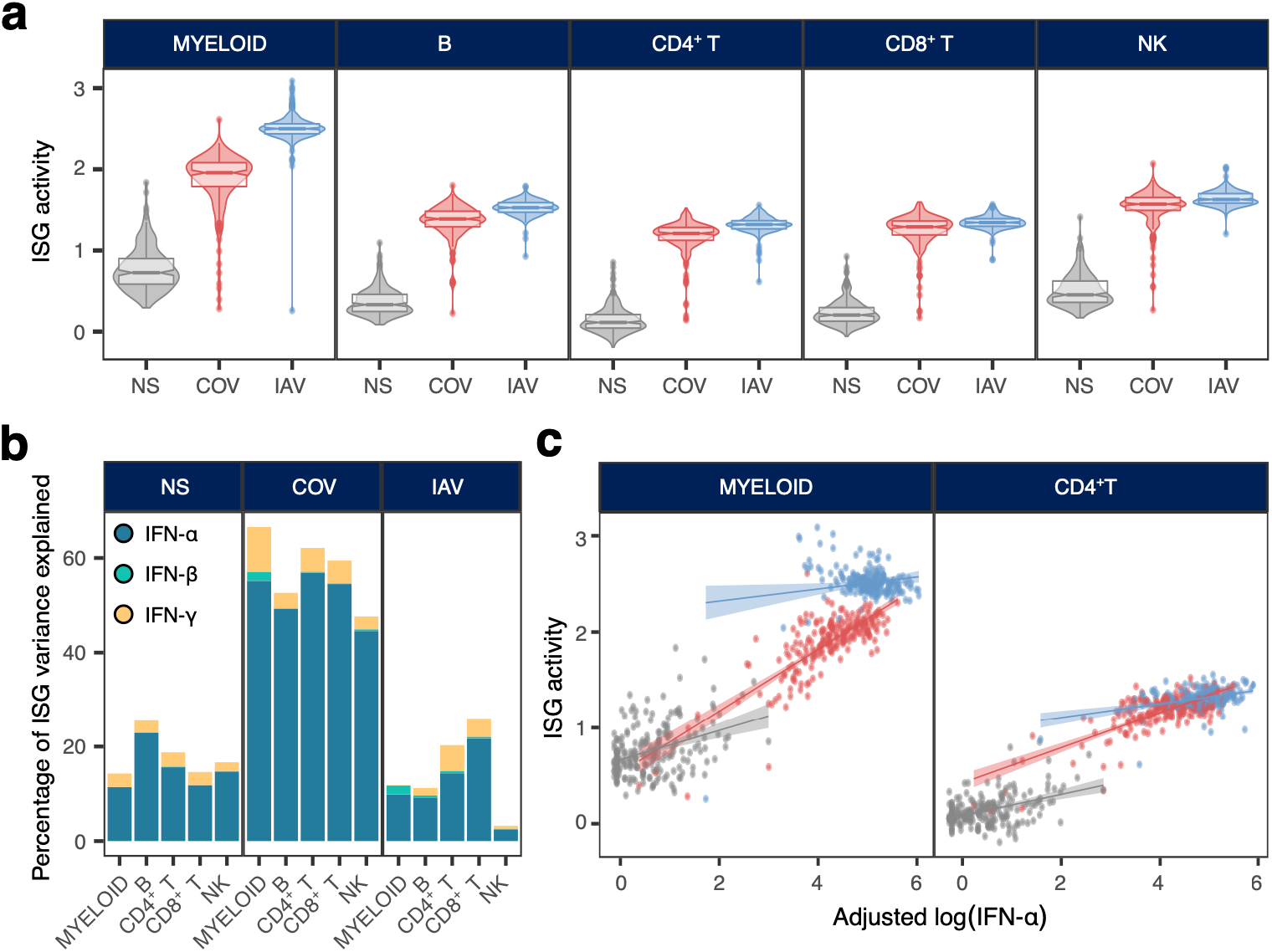
Interferon-stimulated gene responses to SARS-CoV-2 and IAV stimulation. **a,** Distribution of ISG activity in the non-stimulated state and in response to SARS-CoV-2 (COV) and influenza A virus (IAV) across the five major immune lineages. For each lineage and set of stimulation conditions, the violins and boxplots show the distribution of ISG activity scores across all 222 donors (middle line: median; box limits: upper and lower quartiles; whiskers: 1.5× interquartile range; points: outliers). **b**, Proportion of the variance of ISG activity explained by IFN-α, IFN-β and IFN-*γ* in the non-stimulated condition and in response to SARS-CoV-2 and IAV, across the five major immune lineages. **c,** Correlation between levels of IFN-α in the supernatants (measured by SIMOA) and ISG activity in myeloid and lymphoid (CD4^+^ T cells) cells, adjusted for cellular mortality. Each dot represents a sample (donor × condition) and is colored according to its stimulation condition (gray: NS, red: COV and blue: IAV). For each lineage and set of stimulation conditions, regression lines and 95% confidence intervals are shown.

**Supplementary Figure 7.**
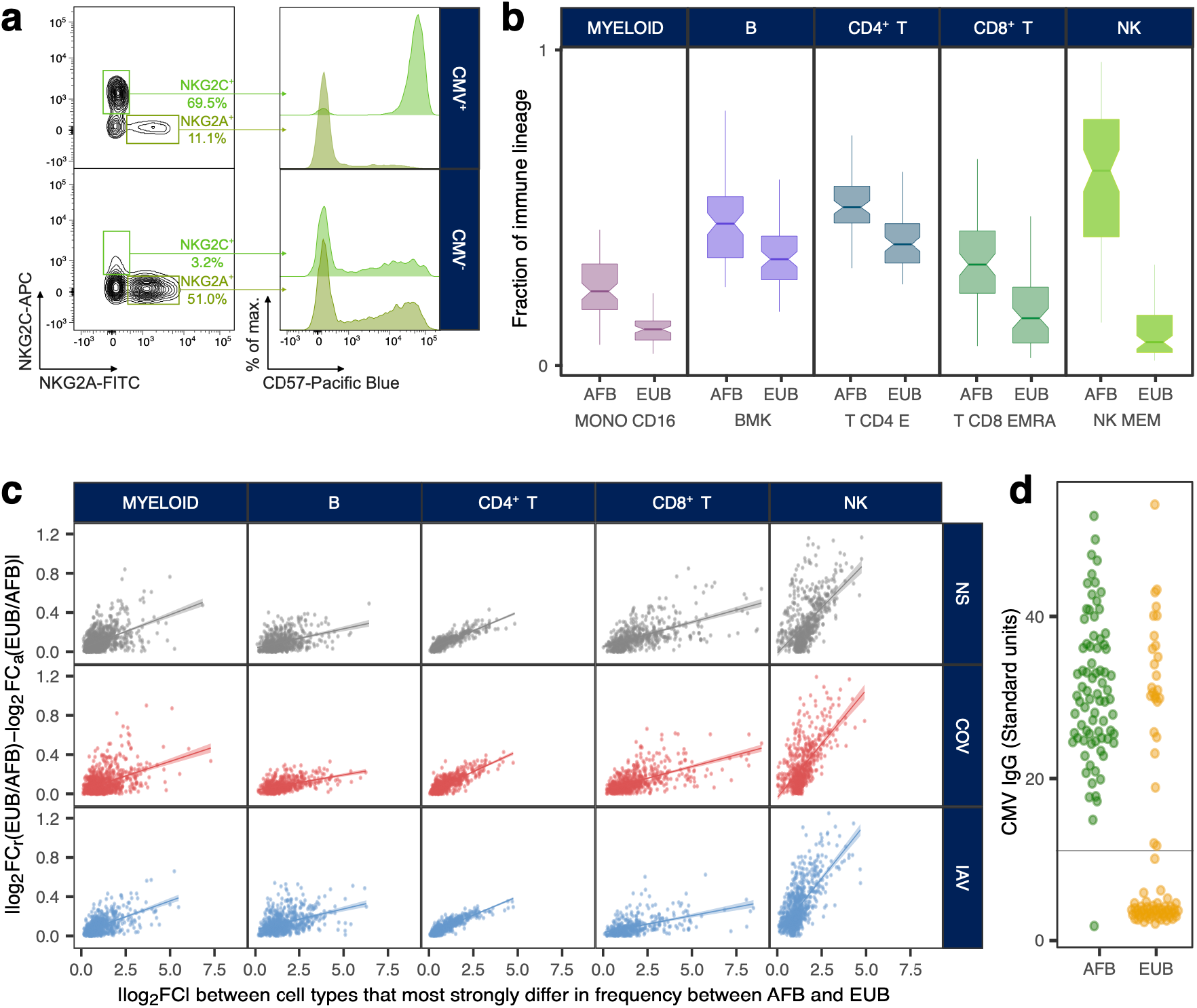
Population cellular heterogeneity and transcriptional responses to viral stimulation. **a,** Validation of the memory-like NK fraction. Flow cytometry data for representative CMV^+^ and CMV^-^ donors, highlighting the higher percentage of memory-like NK cells (NKG2C^+^, NKG2A^-^, CD57^+^) among CMV^+^ donors than among CMV^-^ donors. **b,** Population variation in the percentage of CD16^+^ monocytes, memory lymphocyte subsets and memory-like NK cells. For each major immune lineage, the cell type differing most strongly in frequency between AFB and EUB donors is shown. Boxplots show the distribution of the percentage of the target cell type in the corresponding lineage in each population (middle line: median; box limits: upper and lower quartiles; whiskers: 1.5× interquartile range). **c,** Effect of adjusting for cellular composition on the absolute differences in expression between AFB and EUB donors, as a function of absolute differences in expression between the two cell types differing most in frequency between these populations (Supplementary Table 3b). For each lineage and stimulation condition, the regression line and 95% confidence interval are shown. **d,** Serology assays for CMV across donors according to ancestry. Each dot represents a donor and is colored according to ancestry (AFB: Central Africans, EUB: West Europeans). The gray line represents the detection threshold used to identify a donor as seropositive.

**Supplementary Figure 8.**
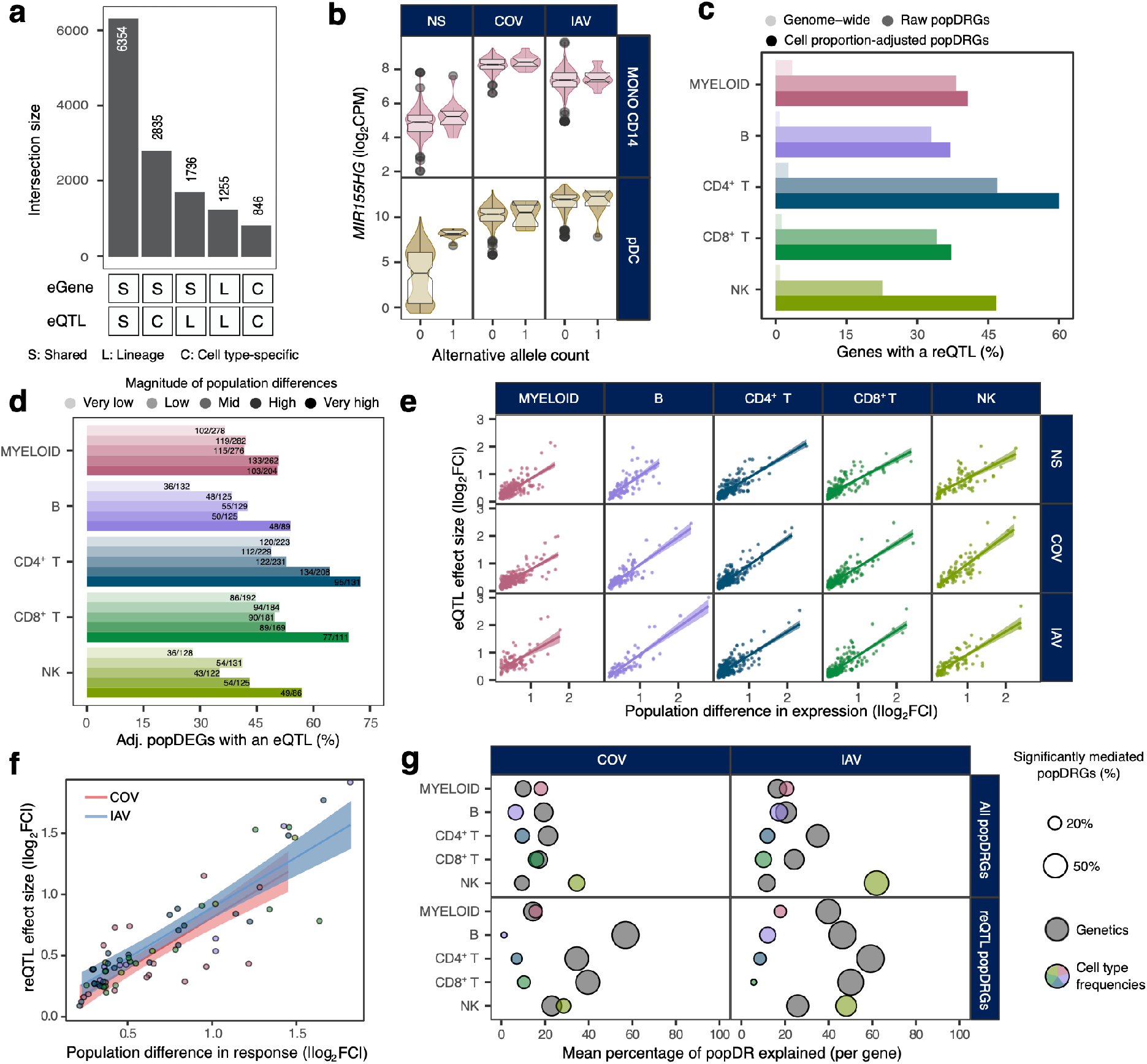
Expression quantitative trait loci mapping and contribution to population differences in response to RNA viruses. **a,** Overlap of eQTLs and eGenes (i.e., genes with an eQTL) detected during the mapping of eQTLs at the immune lineage and cell-type levels. **b,** Example of a pDC-specific eQTL for *MIR155HG. MIR155HG* expression levels in pDCs and CD14^+^ monocytes according to rs114273142 genotype in non-stimulated (NS), SARS-CoV-2-stimulated (COV) and influenza A virus-stimulated (IAV) conditions (middle line: median; box limits: upper and lower quartiles; whiskers: 1.5× interquartile range; points: outliers). **c,** Enrichment in reQTLs among popDRGs. For each lineage, bars indicate the percentage of genes with a significant reQTL, both genome-wide and among the popDRGs identified, before or after adjustment for cell composition (referred to as “adjusted” and “raw” respectively). **d,** Percentage of popDEGs with an eQTL according to the magnitude of differences in expression. In each lineage, popDEGs are assigned to one of five magnitude groups based on quintiles of log_2_fold change between the AFB and EUB populations. For each lineage and magnitude group, the number of popDEGs with an eQTL and the total number of popDEGs are reported. **e,** Relationship between eQTL effect sizes and population differences in expression. **f,** Relationship between reQTL effect sizes and population differences in response to immune stimulation. For each stimulation condition, the regression line is computed jointly across all cell types. **g,** Contribution of genetics and cell composition to population differences in response to stimulation by COV and IAV. For each lineage and stimulation condition, the *x*-axis indicates the mean percentage of population differences in response to stimulation mediated by either genetics or cell composition, across all popDRGs (upper panels) or the set of popDRGs with a significant reQTL (lower panels). The size of the dots reflects the percentage of genes with a significant mediated effect (FDR<1%).

**Supplementary Figure 9.**
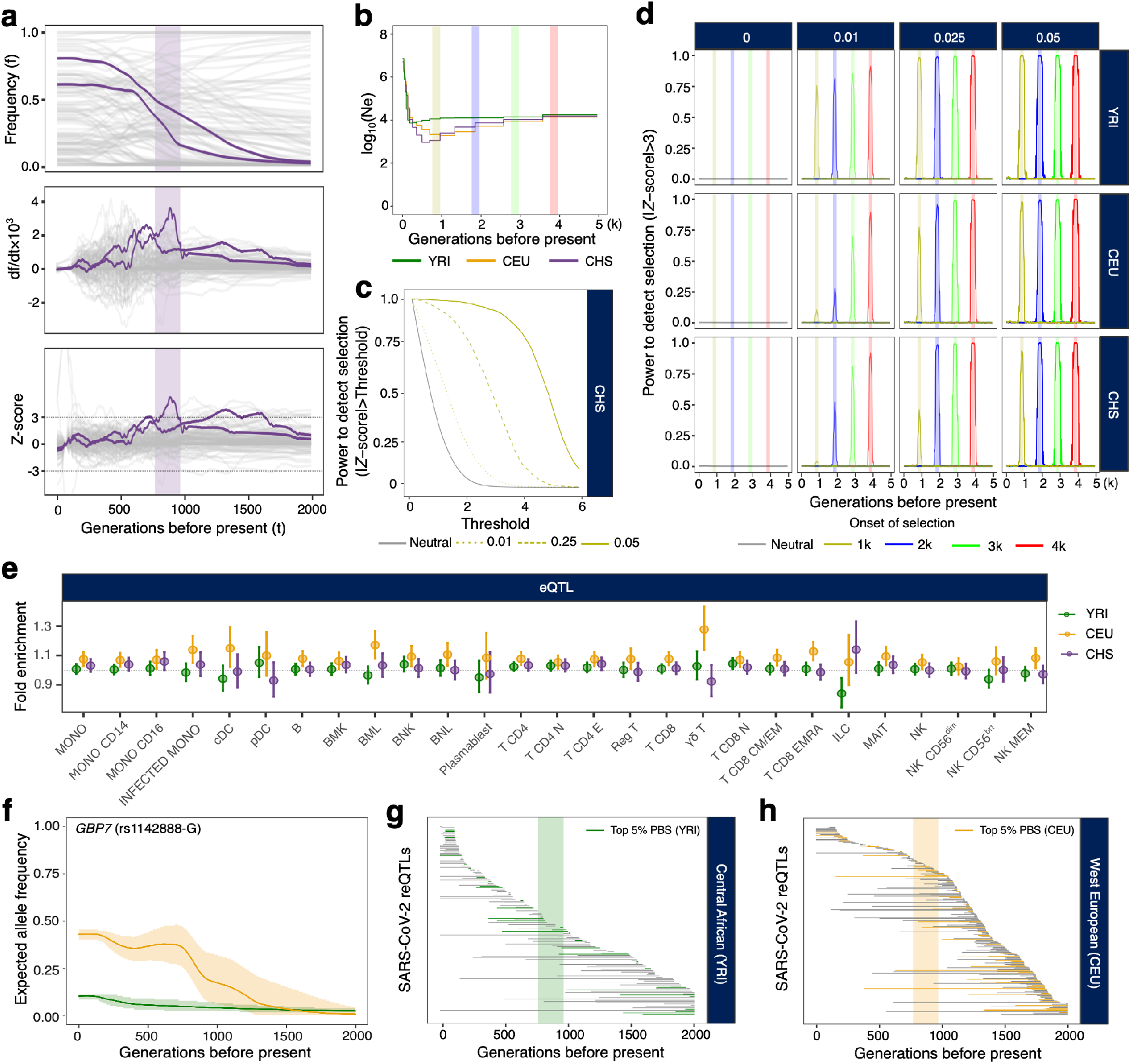
Positive selection signals across time, cell types and populations. **a,** Method for estimating the time of onset of selection from derivative information from allele frequency trajectories. (Top) Allele frequency trajectories in an East Asian population (CHS) across the past 2,000 generations of two SARS-CoV-2 reQTLs (i.e., rs4806787 and rs1028396), affecting the response of *LILRB1* in plasmacytoid dendritic cells and *SIRPA* in CD14^+^ monocytes, respectively. (Middle) Change at each generation (from past to present) of the (smoothed) frequency of the derived allele, normalized for allele frequency. (Bottom) *Z*-score calculated as the normalized derivative, scaled at each generation by the standard deviation of derivatives across all eQTLs. Periods of selection are estimated as the range, in generations, over which the rate of change in the frequency of each allele deviates significantly from expectations under the hypothesis of neutrality (i.e., |*Z*-score| > 3). (Top to bottom) The corresponding allele frequency trajectories, first derivatives and *Z*-scores for 100 random SNPs sampled from the set of all (r)eQTLs detected in this study are shown in gray. **b,** Effective population size and episodes of positive selection over time used in our simulations. Colored lines indicate effective population size (green: YRI, yellow: CEU, purple: CHS); shaded areas indicate positive selection events. **c,** Type I error and power as a function of *Z*-score threshold. Power and type I error are reported for selection occurring 1000-1200 generations ago, for coefficients of selection ranging from 0.01 to 0.05. **d,** Power to detect positive selection at a *Z*-score threshold of 3, as a function of time. Lines are colored according to the date on which selection began (gray for neutral). **e,** Fold enrichments in signals of positive selection (i.e., strong PBS) across the 22 cell types in Central Africans (YRI), West Europeans (CEU) and East Asians (CHS). Fold enrichments were calculated by genome-wide resampling of SNPs matched for minor allele frequency, LD and distance to the nearest gene. Vertical bars indicate 95% confidence intervals. **f,** Allele frequency trajectories over the past 2,000 generations in YRI (green) and CEU (yellow) of the *GBP7* reQTL (rs1142888-G). Shaded areas indicate 95% confidence intervals. **g** and **h,** Estimated period of selection over the past 2,000 generations, for 148 and 279 SARS-CoV-2 reQTLs with significant evidence of natural selection in Central Africans and West Europeans, respectively (max. |*Z*-score| > 3). In both panels, variants presenting strong signals of positive selection (i.e., top 5% for PBS) are shown in color. The transparent rectangle highlights the period between 770 and 970 generations ago (i.e., 21.5-27.2 thousand years ago) associated with polygenic adaptation targeting host coronavirus-interacting proteins. Variants are ordered along the *x*-axis in descending order of time to the onset of natural selection.

**Supplementary Figure 10.**
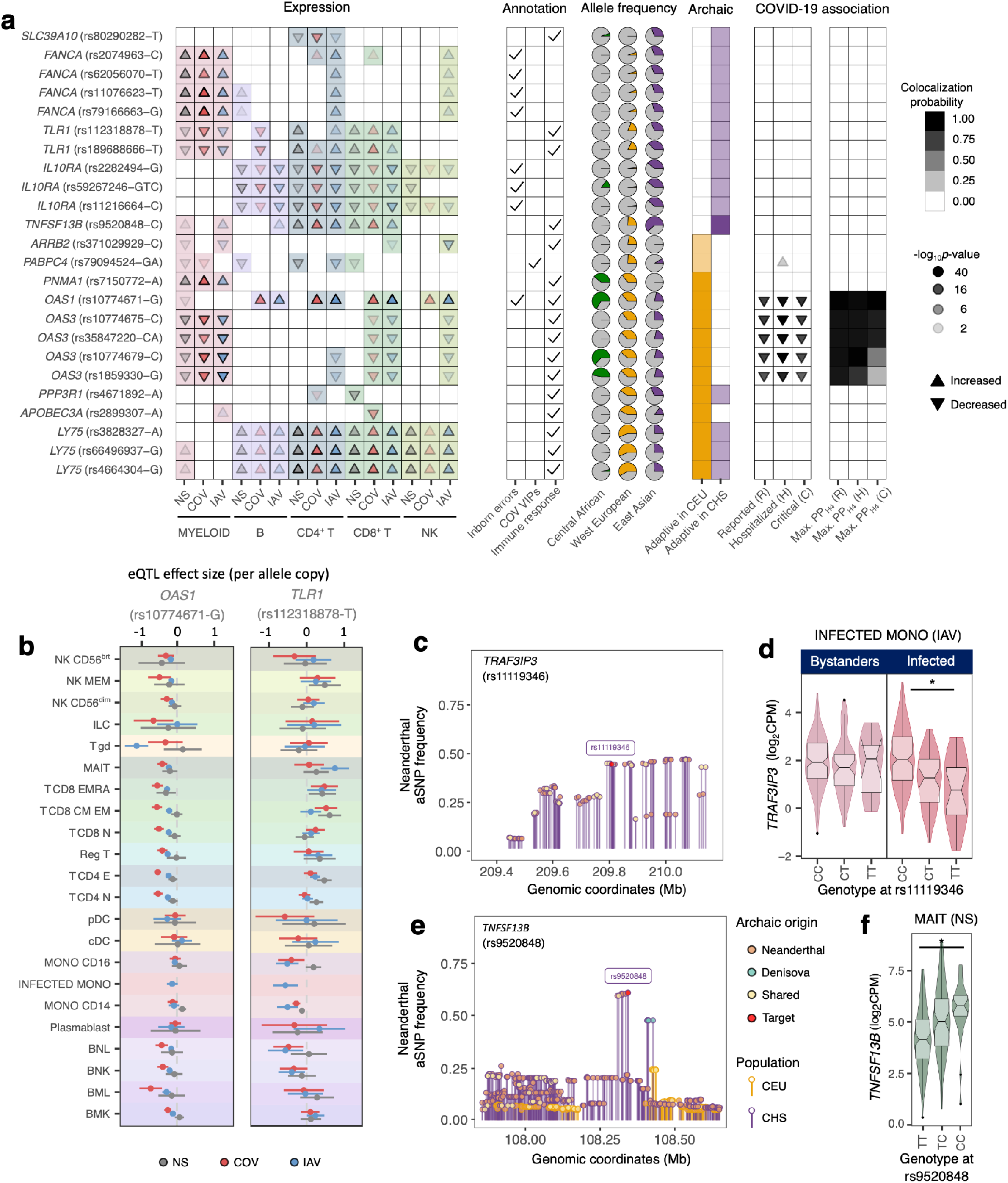
Neanderthal introgression at loci regulating gene expression in different cell types. **a**. Adaptively introgressed eQTLs of host defense genes. From left to right: (i) effects of the introgressed allele on gene expression across immune lineages and stimulation conditions, (ii) clinical and functional annotations of associated genes, (iii) present-day population frequencies of the introgressed alleles, (iv) percentile of archaic allele frequency at the locus (CEU and CHS; dark shades: top 1%, light shades: top 5%, and (v) effects of the target allele on COVID-19 risk (infection, hospitalization, and critical state). Arrows indicate the increase/decrease in gene expression or disease risk with each copy of the introgressed allele. Opacity increases with significance. In the leftmost panel, arrow colors indicate the stimulation condition (gray: NS, red: COV, blue: IAV). For each eQTL, the introgressed allele is defined as the allele segregating with the archaic haplotype in Eurasians. **b**, Effects on gene expression of two loci presenting strong evidence of adaptive introgression (*OAS1, TLR1*). For each locus, eQTL effect size and 95% confidence intervals are shown across the 22 cell types and the three stimulation conditions. **c** and **e**, Frequency and nature of archaic alleles at two introgressed loci (*TNFSF13B* and *TRAF3IP3*) Each dot represents an archaic allele and is colored according to its presence in Vindija Neanderthal (orange), Denisova (green) or both (yellow). The *y*-axis reflects their frequency in CEU (yellow) or CHS (purple) populations. The eQTL index SNP is shown in red. **d**, The Neanderthal-introgressed eQTL at *TRAF3IP3* is apparent only in IAV-infected monocytes, and is not detected in bystander cells (stimulated but not infected). **f**, Effects of the introgressed eQTL at *TNFSF13B* in MAIT cells (i.e., the cell type with the largest effect size). For **d** and **f**, middle line: median; box limits: upper and lower quartiles; whiskers: 1.5× interquartile range; points: outliers.

**Supplementary Figure 11.**
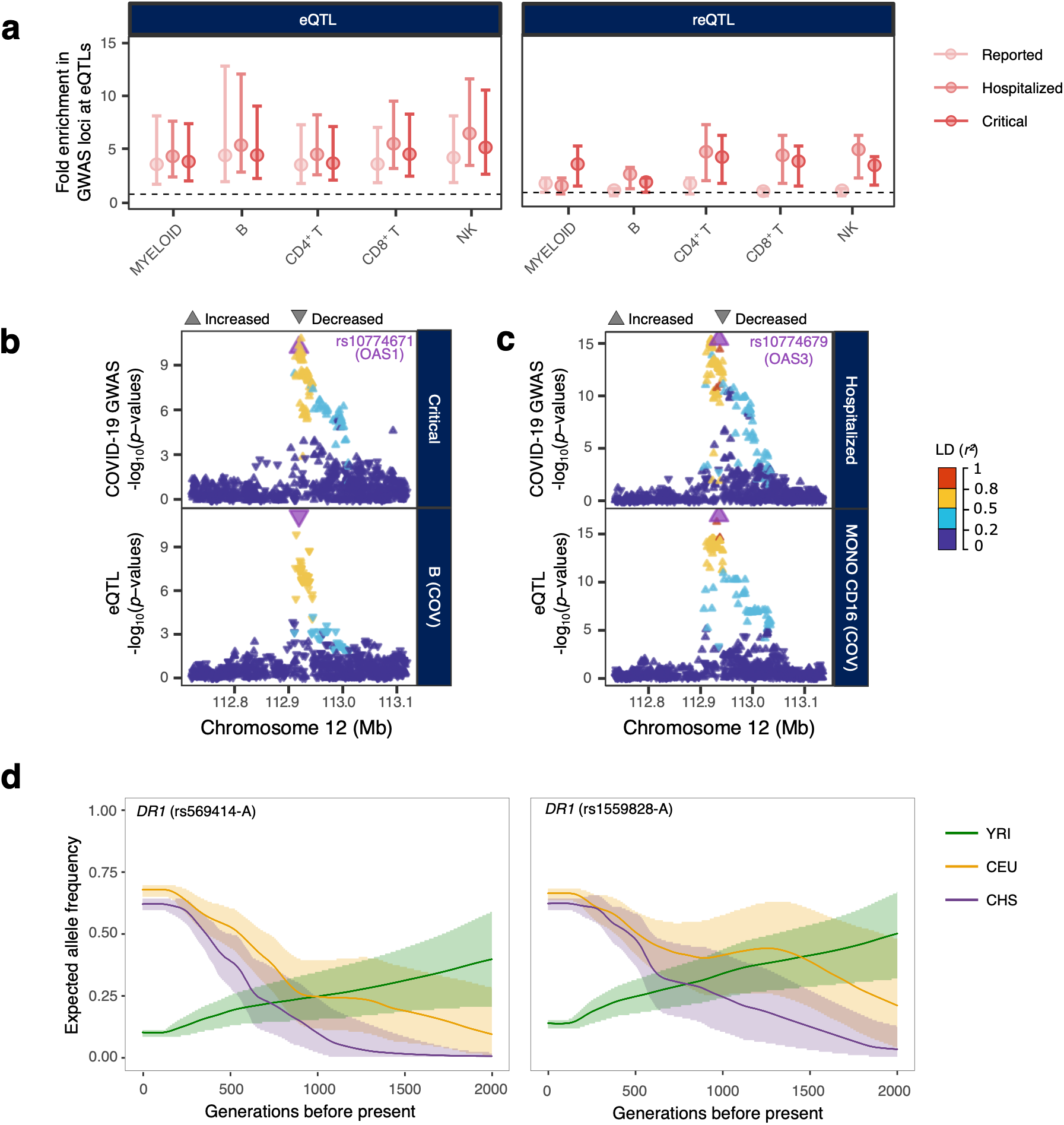
Colocalization of eQTLs and reQTLs with COVID-19-associated loci. **a,** Enrichment in COVID-19-associated loci at eQTLs and reQTLs in each major lineage. For each set of eQTLs and each COVID-19 phenotype, fold enrichment and resampling-based 95% confidence intervals are displayed. **b** and **c,** Colocalization of eQTLs with COVID-19 GWAS hits at the *OAS1-3* locus. For each eQTL, the upper panel shows the log_10_ *p*-value profile for association with COVID-19 phenotypes and the lower panel represents the profile of log_10_ *p*-values for association with expression in a representative cell type. Arrows indicate the direction of the effect at each SNP. The color code reflects LD (*r*^2^) with the consensus SNP, shown in purple, identified by colocalization analysis. **d,** Allele frequency trajectories over the last 2,000 generations in the three populations of two *DR1* eQTLs (rs569414 and rs1559828) that colocalize with COVID-19 severity loci. Shaded areas indicate 95% confidence intervals.

