## Supplementary Notes for "Environmental and genetic drivers of population differences in SARS-CoV-2 immune responses"

### Supplementary Note 1: Single-cell dynamics of the transcriptional responses to SARS-CoV-2 and IAV

To monitor the dynamics of the transcriptional response to SARS-CoV-2, we performed a kinetic single-cell RNA-sequencing experiment on peripheral blood mononuclear cells from a subset of individuals of different ancestry ($n$ = 56 samples, 4 AFB and 4 EUB), following 0, 6 and 24 hours of stimulation with SARS-CoV-2 (BetaCoV/France/GE1973/2020) or IAV (H1N1/PR/8/1934) (Supplementary Fig. 2). After filtering low-quality cells and genetic doublets, we captured 86,177 single-cell transcriptomes, which were assigned to five major cell lineages (myeloid cells, B cells, $CD4^+$ T cells, $CD8^+$ T cells and natural killer (NK) cells) based on per-cluster expression profiles. We found that cell-type heterogeneity drives most of the variation in gene expression (40%), while time point and virus exposure accounted for a joint 22% (Supplementary Fig. 2a, b). Cellular composition remained stable over time, except for myeloid cells, which dropped after 24 hours of IAV stimulation, probably as a result of productive infection and cell death (Supplementary Fig. 2c). Both viruses induced a strong immune response at 6 hours, with 983 differentially expressed genes (DEGs; FDR<0.01, $|\log_2 FC|>0.5$) in response to either of the virus across cell lineages (Supplementary Fig. 2d and Supplementary Table 1a). While IAV productively infected monocytes, we found no evidence for SARS-CoV-2 replication in PBMCs (Supplementary Fig. 2e), consistent with previous findings[1-3].

We observed that transcriptional responses to SARS-CoV-2 and IAV were highly correlated across cell types, with both viruses, especially IAV, strongly inducing interferon-stimulated genes (ISGs) such as *ISG15*, *MX1* or *RSAD2* (Supplementary Fig. 2f). However, monocyte responses were highly heterogeneous, with SARS-CoV-2 inducing a specific transcriptional network enriched in inflammatory-response genes (GO:0006954; odds ratio (OR) >10.2, $p$-value$<7.8\times10^{-14}$) (Supplementary Table 1b, c). Specifically, *IL1B* and *CXCL8* were strongly upregulated in response to SARS-CoV-2 ($\log_2$FC>2.2, $p$-value$< 7.1 \times10^{-20}$) but not IAV ($\log_2$FC<0.01), highlighting the stronger inflammatory potential of SARS-CoV-2 relative to IAV. Protein assays confirmed the weaker induction of type I and II IFNs by SARS-CoV-2 as well as the specific upregulation of inflammatory cytokines, such as IL-1β, IL-6 or TNF-α (Supplementary Fig. 2g and Supplementary Table 1d).

Because monocytes are productively infected by IAV, and not by SARS-CoV-2, we compared the transcriptome of SARS-CoV-2-stimulated monocytes with that of the fraction of non-infected monocytes in the IAV condition. We observed that the inflammatory profile of SARS-CoV-2-stimulated monocytes remained highly significant (GO:0006954; OR> 10.8, $p$-value $< 5.3\times10^{-16}$; Supplementary Table 1e). This observation suggests that differences in the sensing of viral particles, rather than infection itself, account for the increased inflammatory response observed for SARS-CoV-2.

### Supplementary Note 2: Memory-like NK cells are characterized by a strong exhaustion phenotype

Given the reported association of CMV infection with severe COVID-19 (ref.[4]), we investigated how the differences in cellular composition induced by CMV might alter the leukocyte responses to SARS-CoV-2 infection. We found that memory-like NK cells were characterized by a strong exhaustion phenotype. This was characterized by increased basal and stimulated expression of *LAG3* relative to their non-memory counterparts ($log_2$FC>2.1, FDR< $2.2 \times 10^{-12}$) and decreased induction of major effector cytokines such as *IFNG* and *GZMB* upon viral stimulation ($log_2$FC between $CD56^{dim}$ and memory-like NK cells > 1.0 after stimulation by SARS-CoV-2, FDR < $1.5 \times 10^{-6}$; Supplementary Table 3f). Likewise, $CD8^+$ EMRA T cells displayed high expression levels of cytotoxicity genes, such as *GZMB, GNLY* and *NKG7*, relative to central and effector memory T cells, while sharing their ability to elicit a strong inflammatory response relative to naïve $CD8^+$ T cells (i.e., genes that display stronger responses to SARS-CoV-2 in $CD8^+$ EMRA T cells, relative to naïve CD8+ T cells, present a 3.4-fold enrichment in inflammatory genes relative to genome-wide expectations, FDR < $9.0 \times 10^{-4}$).
